# Pharmacological cAMP stimulation via prostaglandin receptors rescues ciliary defects in CEP290-deficient human and mouse models

**DOI:** 10.1101/2023.10.06.561156

**Authors:** France de Malglaive, Iris Barny, Shahd Machroub, Lucas Fares-Taie, Ema Cano, Nicolas Goudin, Tania Attie-Bittach, Josseline Kaplan, Isabelle Perrault, Luis Briseno-Roa, Jean-Michel Rozet, Jean-Philippe Annereau

**Affiliations:** Laboratory of Genetics in Ophthalmology, INSERM UMR1163, Institute of Genetic Diseases, Imagine, Paris, France, 75015; Medetia Pharmaceuticals, Imagine Institute, Paris, France, 75015; Necker Bioimage Analysis Core Facility of the Structure Federative de Recherche Necker, INSERM US24/CNRS UAR3633, Imagine and Paris University, Paris, France, 75015; Laboratory of Embryology and Genetics of Human Malformation, INSERM UMR1163, Institute of Genetic Diseases, Imagine, and Paris University, Paris, France, 75015

## Abstract

The retina’s sensitivity to light depends on the primary cilium of photoreceptors, known as the outer segment (OS). OS defects are a primary cause of inherited retinal dystrophies (IRDs) and can also indicate wider ciliary dysfunctions. One such IRD is Leber congenital amaurosis type 10 (LCA10), which occurs as a monosymptomatic retinal disease or the presenting symptom of syndromic ciliopathies within the Senior-Loken-Joubert-Meckel spectrum. LCA10 patients are born blind but retain dormant photoreceptors for decades, offering the potential for reactivation. AAV-based gene therapies for IRDs cannot accommodate the large *CEP290* gene, prompting the search for innovative treatments.

LCA10 is caused by mutations in *CEP290*, which plays a vital role in ciliation, similar to cAMP signaling. Utilizing fibroblasts displaying variable *CEP290* mutations and associated ciliary defects, we show that exposure to Taprenepag, a specific PGE2 receptor agonist, substantially stimulated cAMP synthesis and consistently improved cilia formation and elongation. We also found that intraperitoneal injection of Taprenepag in mice reached the retina and slowed retinal degeneration, promoting OS formation, and improving light sensitivity in a *Cep290* mutant mouse. These findings demonstrate the potential of Taprenepag to treat the CEP290-related visual dysfunction and suggest evaluation for other CEP290-related organ issues and ciliopathies.

## INTRODUCTION

Inherited Retinal Dystrophies (IRDs) encompass a diverse group of disorders characterized by the degeneration of rod and/or cone photoreceptors in a diffuse or generalized manner, leading to variable degrees of visual impairment. IRDs collectively affect approximately 1 in 3000 individuals ^1^ and are a primary cause of severe visual deficiency, especially among children.

In addition to exhibiting considerable clinical diversity, IRDs display extensive genetic variability (> 300 genes) ^2^, and a wide range of pathophysiological mechanisms, encompassing both retina-specific and broader cellular pathways ^2^. Ciliary dysfunction represents a prominent mechanism in IRDs, referred to as retinal ciliopathies; this prominence can be attributed to the fact that photoreceptor cells sense light through a modified primary cilium referred to as the outer segment (OS), essential for capturing and transducing incoming light into electrical signals, the foundation of visual perception ^3,4^.

Numerous genes playing pivotal roles in primary cilium biogenesis, maintenance, and function, have been associated to retinal ciliopathies that may manifest as non-syndromic diseases or as overlapping syndromes that involve principally the central nervous system, kidney, and bones together with the retina. Among them, *CEP290* stands out as a major contributor to Leber congenital amaurosis (LCA), which represents the earliest and most severe form of IRDs, responsible for blindness or profound visual deficiency at or near birth. This disease is a leading cause of childhood blindness, affecting approximately 20% of children enrolled in schools for the blinds ^5^. To date, over 20 genetic subtypes of LCA have been reported ^6^, with LCA10, attributed to mutations in the *CEP290* gene, being the most prominent subtype (20% of all LCA cases) ^6^. Interestingly, *CEP290* also plays a substantial role in LCA forms with neuro-renal features of the syndromic Senior-Loken-Joubert-Meckel spectrum ^7^.

CEP290 has demonstrated essential functional and structural roles in cilia formation. Consistently, studies using CEP290 mouse models and patient-derived induced pluripotent stem cell (iPSC)-derived retinal organoids have revealed normally developed retinas but absent or rudimentary outer segments (OS), resulting in an inadequate response to light stimulation ^8–11^. This finding aligns with the profound visual impairment experienced by affected children at or around birth. However, high-resolution retinal imaging and functional magnetic resonance neuroimaging demonstrate that individuals with *CEP290* mutations retain dormant photoreceptors in their central retina for over two to three decades while maintaining intact visual pathways ^12,13^. These findings offer promising prospects for therapies aimed at preserving and activating these dormant cells to restore light perception.

Delivering a functional copy of the mutated gene using adeno-associated viral (AAV) vectors into the retina has demonstrated significant promise in preclinical and clinical studies. Notably, a product has successfully reached the market for treating RPE65-associated LCA (LCA2), marking a significant milestone in the field ^14,15^. However, LCA10 poses a distinct challenge due to the size of the *CEP290* cDNA (7.2 Kb), exceeding conventional AAV vector capacities (< 5 Kb). Consequently, alternative strategies, such as antisense oligonucleotide (ASO) ^1,16–19^ and CRISPR-Cas9-mediated therapies ^20,21^, have advanced to correct a highly recurrent deep intronic mutation responsible for LCA10, but their specificity limits their impact to the mutation in question.

In our study, we focused on a small drug mutation-agnostic therapeutic approach. We report that Taprenepag, a specific agonist of prostaglandin E_2_ (PGE2) receptor EP_2_ subtype (EP_2_), can enhance cilia formation in fibroblasts with *CEP290* mutations and related cilia formation defects of varying severity. This enhancement is achieved by promoting cAMP synthesis, which can be achieved with low dose such as 0.2µM. Additionally, our research demonstrates that repeated intraperitoneal injections of Taprenepag, tested in a *Cep290* mutant mouse model characterized by rapid and severe retinal degeneration, not only prove highly safe but also effectively slow disease progression and improve retinal sensitivity to light by substantially promoting OS formation and increasing photopigments availability. These findings lay the groundwork for potential preclinical studies and the exploration of Taprenepag as a promising therapeutic intervention for individuals with CEP290-related retinal disorders.

## RESULTS

### CEP290 mutant fibroblasts manifest variable cilia formation defects

We cultured fibroblasts carrying *CEP290* mutations of varying severity as well as control in a serum-deprived medium to stimulate cilia formation. Subsequently, we quantified the proportion of ciliated cells and assessed cilium sizes across the cell lines.

We found significant reductions in the number of cells with primary cilia in all mutant fibroblasts compared to controls. The reduction was statistically significant in both the E14/E32 and E36 lines, which carry very severe and severe *CEP290* mutations, but not in the I26 line, which carries a hypomorphic *CEP290* mutation. Specifically, we found that only 9.2% of E14/E32 cells, 39.6% of E36 cells, and 63.8% of I26 cells had primary cilia, whereas controls had a significantly higher average percentage of ciliated cells at 66.8% (1-way ANOVA with Dunnett multiple comparison; n=3; p<0.0001, p<0.0001 and p=0.8940 respectively; **Figure 1A and B**).

**Figure 1:**
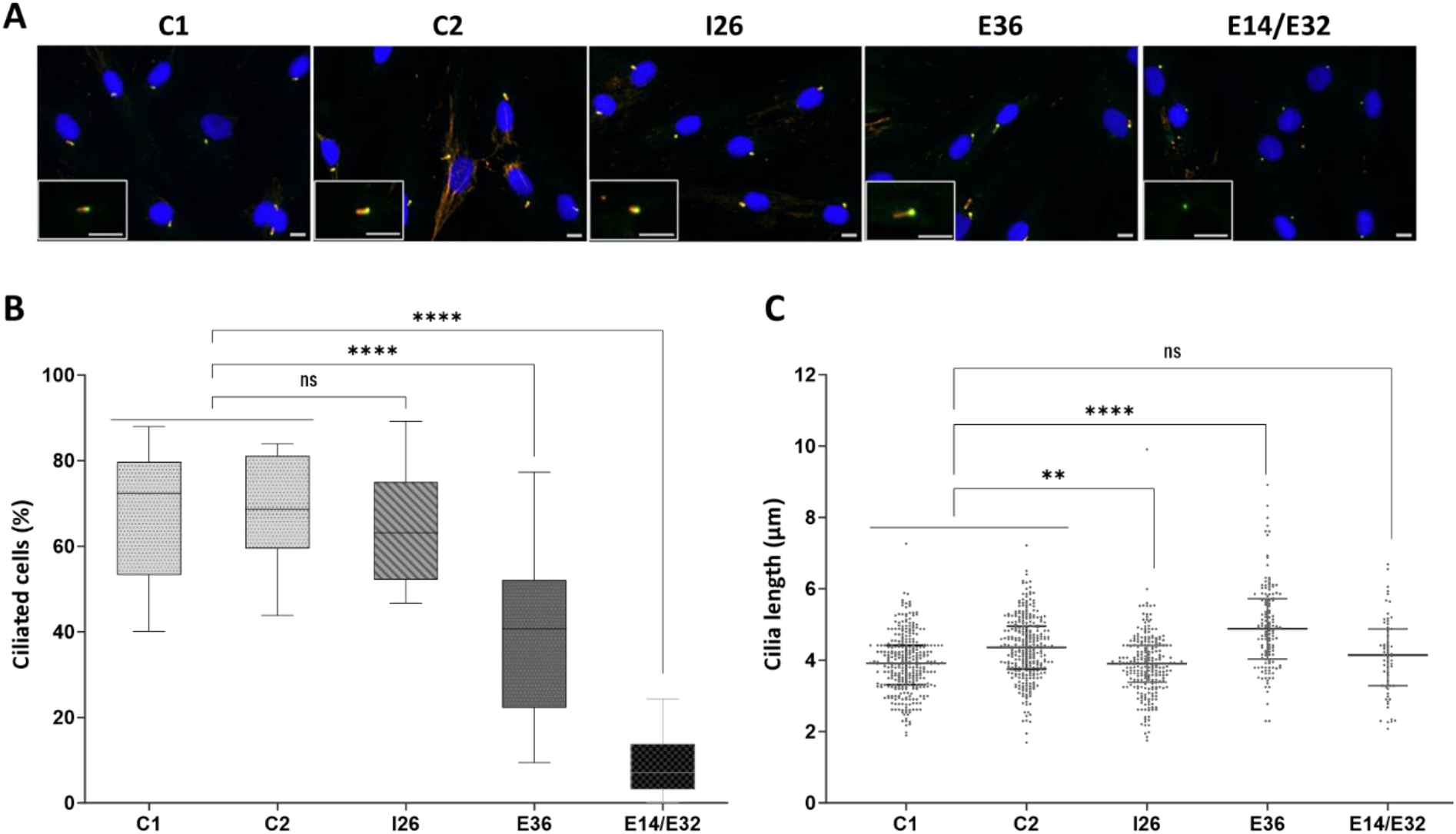
Ciliogenesis in control and mutant fibroblasts. **A.** Representative images displaying cilia in control (C1, C2) and mutant (I26, E36, E14/E32) primary fibroblasts under serum starvation. Cilia are marked by ARL13B (red) labeling, basal bodies by Gamma-tubulin (green), and nuclei by DAPI (blue). Scale bars are 20µm for the main images and 10µm for the inserts. **B.** Quantitative assessment of the percentage of ciliated cells. Data is presented in a boxplot format, with the horizontal line within the box representing the median. The upper and lower boundaries of the box denote the 25th (lower quartile Q1) and 75th (upper quartile Q3) percentiles, respectively. The whiskers extending from the box represent the 10th and 90th percentiles. **C.** Quantitative analysis of the average cilia length. Each data point corresponds to a single cilium. Bars indicate the median with Q1 and Q3 values. Data in panels B and C were obtained from three independent replicates, with a sample size (N) exceeding 250 cells. Statistical analysis was performed using an ordinary one-way ANOVA followed by Dunnett’s multiple comparison test. Significant p-values are denoted as follows: **** for p < 0.0001 and ** for p < 0.01. “ns” indicates non-significance.

Additionally, ciliary lengths in mutant cells consistently exhibited significant abnormalities, with I26 cells having shorter cilia and E36 cells having longer cilia when compared to controls (1-way ANOVA with Dunnett multiple comparison; n>250 cells; p=0.0013, p<0.0001 respectively; **Figure 1C**).

In summary, these results collectively highlight variable but noteworthy cilia formation defects in the three fibroblast lines with diverse *CEP290* mutations. These findings are consistent with the evaluation of compounds aimed at addressing cilia formation issues.

### *PTGER_2_* encoding EP_2_ is expressed in fibroblasts and impacted by Taprenepag exposure

Taprenepag is a potent and selective PGE_2_ agonist with a very high selectivity for the human EP_2_ receptor over EP_1,_ EP_3,_ EP_4_ (approximately 270 folds ^22^). Before evaluating Taprenepag impact on cAMP synthesis in fibroblasts, we examined the baseline expression of the *PTGER_1_, PTGER_2_, PTGER_3_* and *PTGER_4_* genes in human primary fibroblasts using end-point RT-PCR analysis. In controls, we readily detected *PTGER_1_* and *PTGER_4_* mRNA, while *PTGER_3_* mRNA was not detectable (**Figure S1**). Additionally, *PTGER_2_* mRNA was detectable in both control cells and mutant cells carrying various *CEP290* genotypes (**Figure 2A and Figure S1**).

**Figure 2:**
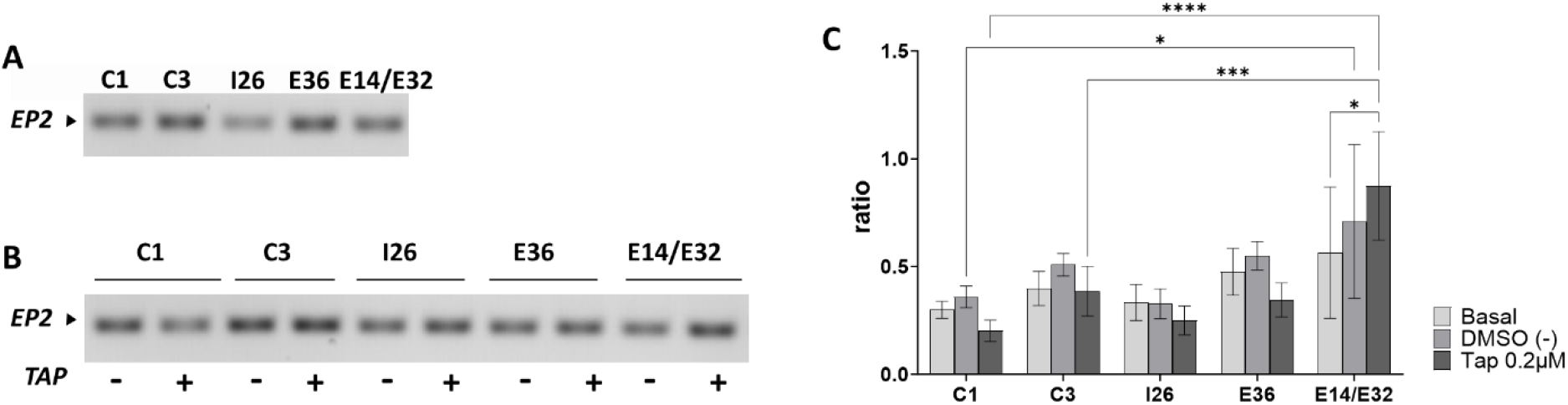
*PTGER_2_* mRNA expression in control and mutant fibroblasts carrying various *CEP290* mutations under different conditions. **A.** Detection of *PTGER_2_ (EP_2_)* mRNA in untreated control (C1, C3) and mutant (I26, E36, E14/E32) fibroblasts under serum starvation. **B.** Detection of *EP_2_* mRNA in control (C1, C3) and mutant (I26, E36, E14/E32) fibroblasts under serum starvation after 24 hours of exposure to either the solvent DMSO (-) or 0.2µM Taprenepag (+). **C.** Relative expression of *PTGER_2_* mRNA in control (C1, C3) and mutant (I26, E36, E14/E32) fibroblasts under serum starvation conditions with no treatment, or treatment with DMSO or Taprenepag in DMSO. The analysis was conducted using quantitative RT-PCR with *GUSB* and *RPLP0* genes as reference genes. Bars represent the mean ± standard deviation from ≥ 2 independent replicates. Statistical analysis involved an ordinary two-way ANOVA followed by a Tukey’s multiple comparison test. Significant p-values are denoted as follows: **** for p < 0.0001, *** for p < 0.001, and * for p < 0.05.

We subsequently investigated whether a 48h exposure to Taprenepag at a concentration of 0.2µM (10 times the EC50 reported in HEK293 cells ^22^), or vehicle (DMSO) in a serum-free culture medium influenced gene expression using RT-PCR and RT-qPCR analyses. When cells from I26, E36, and control lines were exposed to DMSO, a slight increase in *PTGER_2_* mRNA content compared to basal levels was observed. Conversely, a more pronounced decrease in *PTGER_2_* mRNA content was consistently observed across all these cell lines upon exposure to Taprenepag in DMSO. While statistically insignificant, these trends were uniform among all cell lines, with minimal variations between them **(Figure 2B and C**), suggesting potential biological relevance. Specifically, these observations point to a dampening effect on *PTGER_2_* mRNA expression induced by Taprenepag, which may have been offset to some extent by the opposite effect of DMSO. In the E14/E32 line, we observed elevated basal levels of *PTGER_2_* mRNA compared to the other cell lines. Exposure to DMSO alone resulted in a slight increase in *PTGER_2_* mRNA content, similar to the effect observed in other cell lines. However, upon exposure to Taprenepag in DMSO, the E14/E32 line exhibited a notably more substantial increase in *PTGER_2_* mRNA content, in stark contrast to the reduction observed in the other lines. Consequently, a statistically significant elevation in *PTGER_2_* content was noted in the E14/E32 line relative to the other lines following Taprenepag treatment. This intriguing observation was accompanied by higher variability among experimental replicates in the E14/E32 line compared to the other lines (**Figure 2C**).

Collectively, these results demonstrate that the *PTGER_2_* gene is expressed in cultured fibroblasts and impacted by Taprenepag in a similar manner in all cell lines except for the E14/E32 line.

### Taprenepag stimulates cAMP synthesis in mutant and control fibroblasts

To evaluate the effect of Taprenepag on cAMP synthesis in mutant and control fibroblasts, which exhibited varying cilia formation phenotypes, the cells were exposed to 0.2μM and 2μM of the compound or DMSO (sham) in a serum-free culture medium. As a positive control, we exposed the same cell lines to Forskolin, a widely used cell-permeable adenylyl cyclase allosteric agonist, at concentrations previously reported to enhance ciliogenesis (25µM ^23^), as well as a lower dose of one-tenth (2.5µM).

After a 10-minute exposure to 0.2µM of Taprenepag, the intracellular cAMP concentrations increased by 32 to 68% in all cell lines compared to the sham treatment (2-way ANOVA with Dunnet’s comparison; n=3; p<0.01). Treatment with the 2µM dose resulted in an additional, yet non-significant, increase in cAMP concentration, leading to a few-fold rise in all cell lines except the E14/E32 cells. In E14/E32 cells, the fold change in cAMP concentration was consistent upon exposure to 0.2 and 2µM (**Figure 3**).

**Figure 3:**
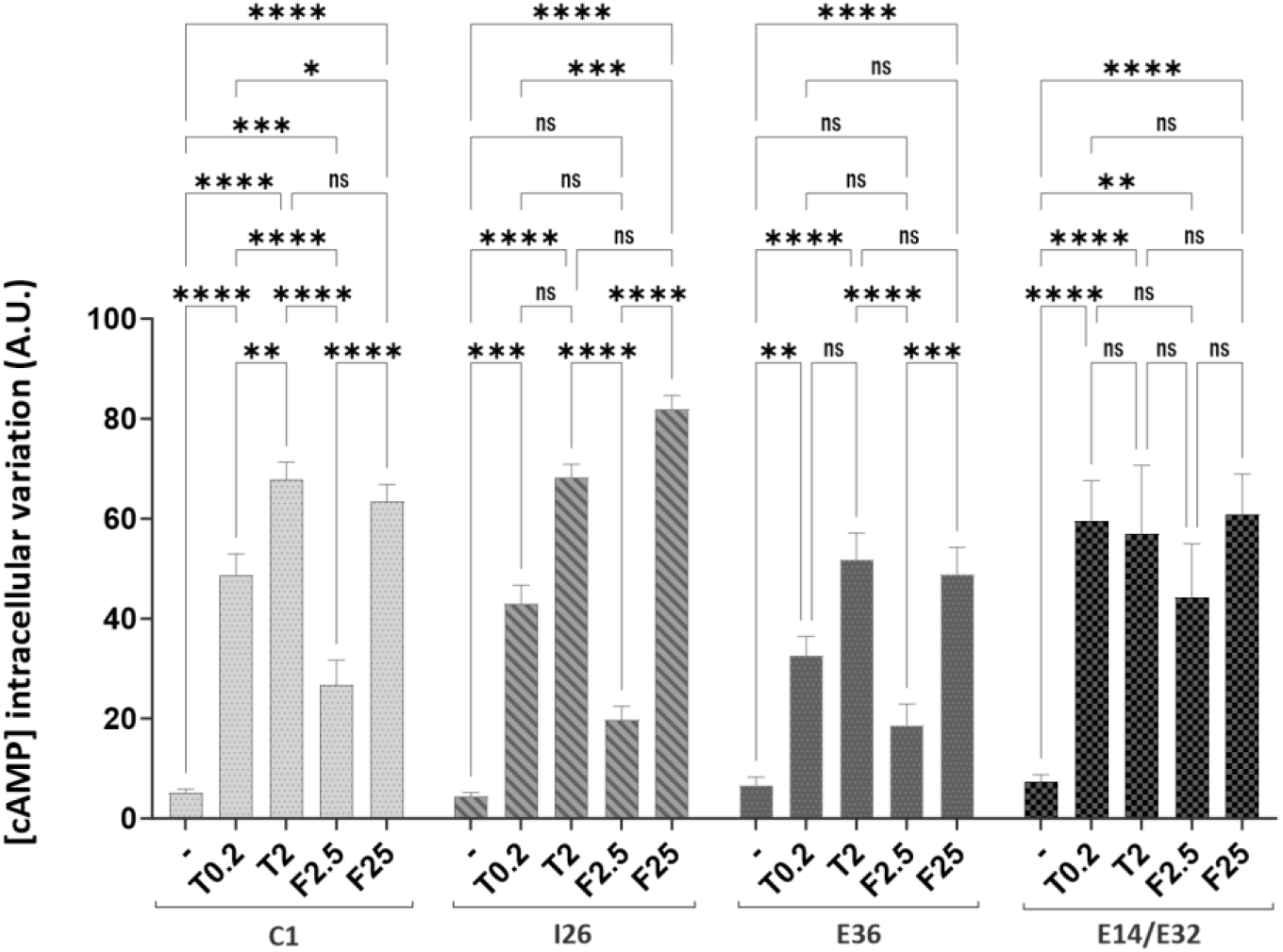
cAMP induction by Forskolin and Taprenepag in control and mutant fibroblasts. This figure illustrates the variation in intracellular cAMP concentration in both control (C1) and mutant (I26, E36, E14/E32) cells after different treatments, including DMSO (-), 0.2µM Taprenepag (T0.2), 2µM Taprenepag (T2), 2.5µM Forskolin (F2.5), and 25µM Forskolin (F25). The bars represent the mean values ± SEM from three independent experimental replicates. The units are represented as arbitrary units (A.U.). Statistical analysis involved an ordinary two-way ANOVA followed by Dunnett’s multiple comparison test. Significant p-values are denoted as follows: **** for p < 0.0001, *** for p < 0.001, ** for p < 0.01, * for p < 0.05, and ns for not significant.

Treatment with 2.5µM of Forskolin increased the intracellular cAMP levels in all cell lines but to a lesser extent than 0.2µM Taprenepag, yet not significant. When the cells were exposed to 25µM Forskolin, there was a notable elevation in cAMP levels, which reached levels similar to those observed following treatment with a 2µM dose of Taprenepag, without exceeding them (2-way ANOVA with Dunnet’s comparison; n=3; non-significant; **Figure 3).**

Collectively, these findings demonstrate that exposure to Taprenepag strongly increases cAMP synthesis. The level of stimulation is of greater potency than the pharmacological positive control, Forskolin, in stimulating total cAMP cell synthesis. Additionally, all cell lines responded to the drugs in a dose-dependent manner, apart from the E14/E32 line upon Taprenepag treatment. However, the E14/E32 line response to the 0.2μM treatment was as high as in other cell lines, supporting further analysis.

### Taprenepag enhances ciliogenesis in both mutant and controls fibroblasts

To assess Taprenepag’s capability in promoting cilia formation, we exposed both mutant and control fibroblasts to a concentration of 0.2µM of the compound, while using DMSO as a sham control, and Forskolin (2.5µM and 25µM) as positive control. These exposures took place in a serum-free culture medium to promote cilia formation. Treatment efficacy was measured by comparing the proportion of ciliated cells and the ciliary lengths across these different experimental conditions.

Exposure to 0.2µM Taprenepag resulted in a significant increase in the proportion of cells harboring a primary cilium in E14/E32 and E36 fibroblasts, which initially exhibited the most severe cilia depletion at the basal level. In the I26 and control groups, we observed a consistent trend towards an increase (specific values and p-values can be found in **Figure 4B**).

**Figure 4:**
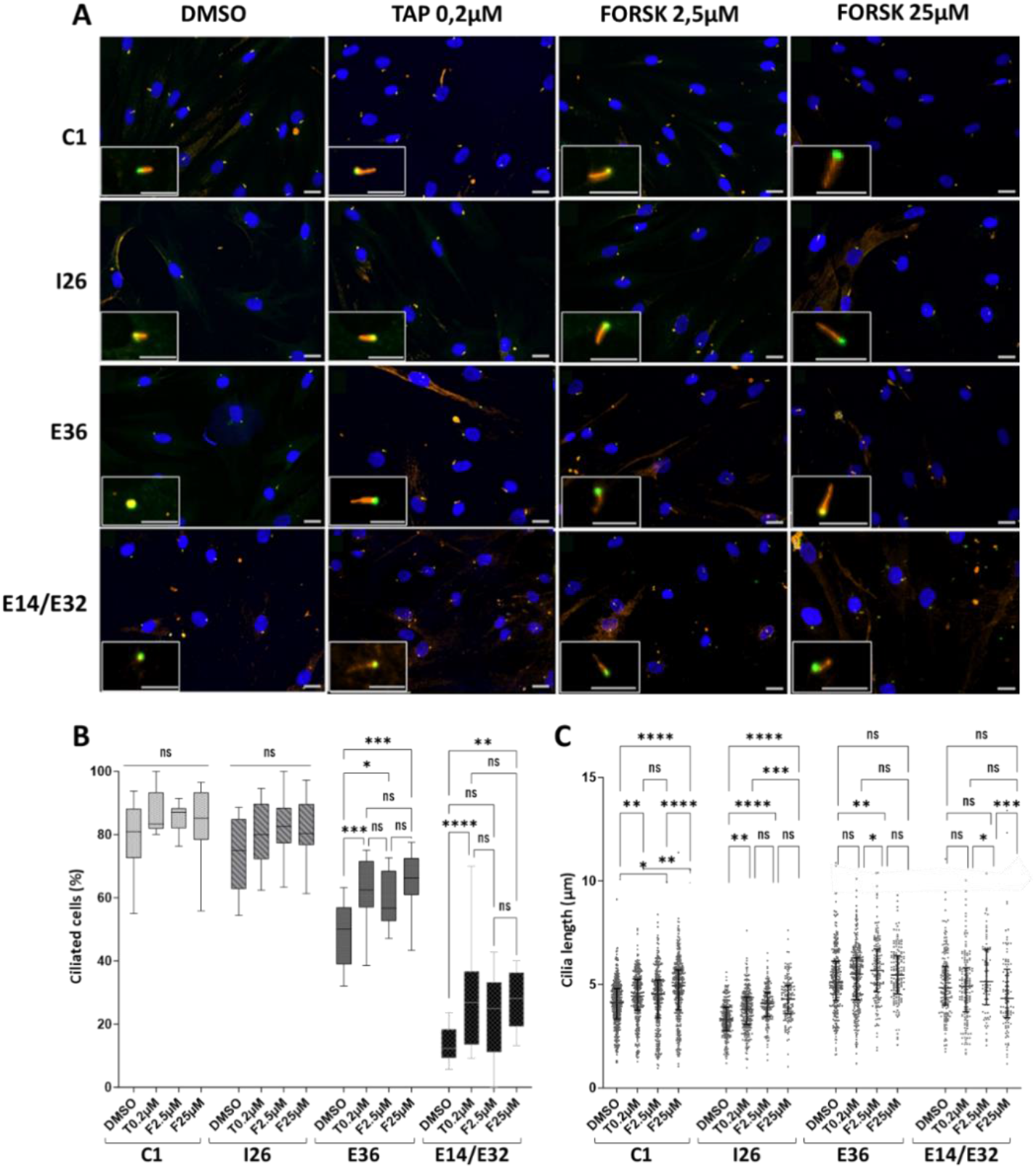
Taprenepag and Forskolin effects on ciliary phenotypes in control and mutant fibroblasts. **A.** This panel displays representative images of cilia in control (C1) and mutant (I26, E36, E14/E32) fibroblasts. These cells were exposed to either DMSO, 0.2µM Taprenepag (TAP), or 2.5µM and 25µM Forskolin (FORSK). Red labeling represents ARL13B, marking the ciliary membrane, green labeling corresponds to Gamma-tubulin, indicating the basal body, and blue labeling is DAPI, marking the nuclei. Scale bars are provided for reference, with 20µm for the main images and 10µm for the inserts. **B.** Quantification of the percentage of ciliated cells in both control and mutant fibroblasts following exposure to DMSO, 0.2µM Taprenepag (T0.2µM), or 2.5µM and 25µM Forskolin (F2.5µM and F25µM, respectively). The data are presented in a boxplot graphical view. The horizontal line within the box represents the median, while the upper and lower limits of the box represent the 25th (Q1) and 75th (Q3) percentile values. Whiskers extending from the box represent the 10th and 90th percentiles. **C.** Quantification of the average cilia length after exposure to DMSO, 0.2µM Taprenepag, or 2.5µM and 25µM Forskolin. Each data point corresponds to a single cilium, with bars indicating the median values along with Q1 and Q3. Data presented in panel B and C were collected from three independent replicates, involving more than 250 cells. Statistical analysis was performed using an ordinary one-way ANOVA followed by Tukey’s multiple comparison test. Significant p-values are represented as follows: **** for p < 0.0001, *** for p < 0.001, ** for p < 0.01, * for p < 0.05, and ns for not significant.

Regarding cilia length, we observed a significant mean elongation of 0.5 to 1µm in the E36, I26, and control fibroblasts (**Figure 4A and C**). Notably, in the E14/E32 line, which initially displayed a wide variety of ciliary lengths, the Taprenepag treatment did not reduce the overall variability but increased the number of cilia with lengths within the control range (**Figure 4C**, **Figure S2**).

Previously, we reported that the I26 and E36 lines expressed minimal amounts of wildtype and minimally shortened CEP290 proteins, respectively ^1,17,24^. To investigate whether the enhancement of cilia formation could be attributed to an increased abundance of CEP290 following Taprenepag exposure, we analyzed the CEP290 protein content in cells exposed to DMSO and Taprenepag in DMSO, respectively, using Western blot analysis. We observed consistent levels of CEP290 protein in all cell lines and under various experimental conditions, indicating that the improvement in cilia formation is unlikely to be attributed to increased expression or extended half-life of CEP290 in the cells (**Figure S3**).

Exposure of serum-starved fibroblasts to Forskolin increased the proportion of ciliated cells in a lower magnitude than 0.2µM Taprenepag. While a slight tendency towards an increased effect was noted in the E14/E32 and E36 cell lines when exposed to 25µM of Forskolin, the fold change in the abundance of ciliated cells remained consistent when exposed to both 2.5 and 25µM in the I26 and control lines (**Figure 4**).

These results illustrate that Taprenepag enhances cilia formation independently of CEP290 at nanomolar concentrations, while Forskolin requires micromolar concentrations, thereby supporting the notion of Taprenepag’s higher potency.

### Absence of synergistic ciliogenesis enhancement with non-selective EP-receptor activation by Alprostadil further supports the key role of EP_2_

To provide additional confirmation that Taprenepag’s effect in fibroblasts is indeed mediated through the EP_2_ receptor, we investigated the impact of Alprostadil (PGE_1_). Alprostadil has affinity constants of 36 nM for EP_1_, 10 nM for EP_2_, 2.1 nM for EP_4_ and 1.1 nM for EP_3_ ^25^. When we exposed both mutant and control fibroblasts to an active concentration of Alprostadil (0.2µM), which targets all EP receptors except for EP_3_ (mRNA undetected in fibroblasts), we observed a consistent increase in cilia length across all cell lines (**Figures 5 and S4**). This effect mirrored what we observed with a 0.2µM concentration of Taprenepag, known to be active for EP_2_ but inactive for EP_1_ and EP_4_ (**Figures 5 and S4**). The comparison between the two compounds was statistically insignificant (2-way ANOVA with Tukey’s multiple comparison; n=3; p=n.s.), suggesting a similar impact of Alprostadil and Taprenepag on cilia length in this context. The lack of a synergistic effect observed upon exposure to Alprostadil suggests that EP_1_ and EP_4_ likely have minimal or no involvement in cilia elongation in fibroblasts (**Figure S1**).

**Figure 5:**
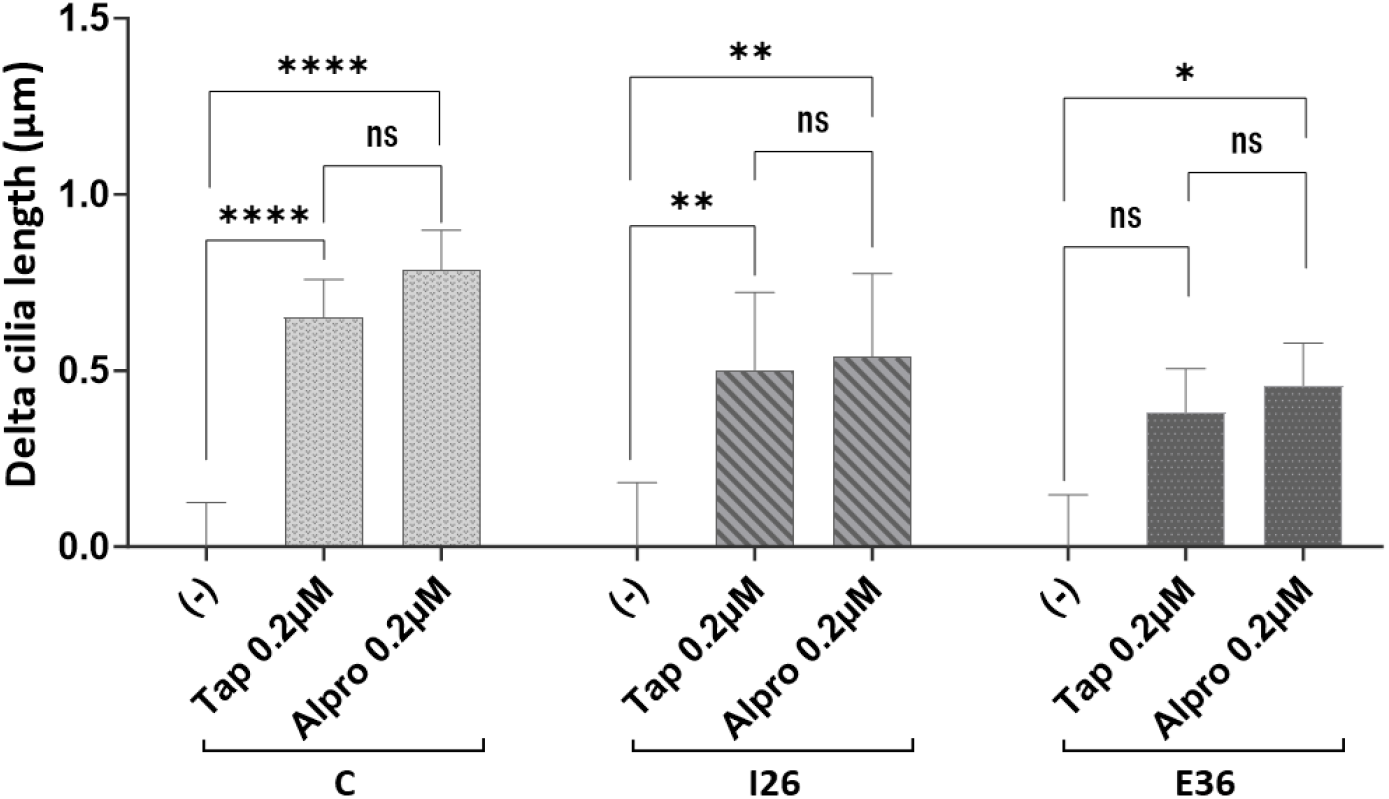
Comparative effect of Taprenepag and Alprostadil on ciliary lengths in control and mutant fibroblasts. This figure illustrates the ciliary phenotype of control (C: pooled values of C1 and C3) and mutant (I26, E36) fibroblasts after 24 hours of serum starvation under “high-content” conditions (refer to Material & Methods for details). The cells underwent serum starvation, followed by a 24-hour exposure to either DMSO (-), 0.2µM Taprenepag, or 0.2µM Alprostadil, both diluted in DMSO. The values represented in the figure depict the difference between the average cilia length of cells treated with the drugs and the average cilia length of cells exposed to DMSO only, as determined by automatic analysis. Each bar corresponds to the mean ± SEM from three independent replicates. Statistical analysis was conducted using an ordinary two-way ANOVA, followed by a Tukey’s multiple comparison test for each cell line. Significant p-values are denoted as follows: **** for p < 0.0001, ** for p < 0.01, * for p < 0.05, and ns for not significant.

### cAMP analogs in the culture medium can enhance ciliation in mutant and control fibroblasts

We explored the potential of cell-permeable and stable cAMP analogs in the culture medium to promote cilia formation in I26, E36, and control fibroblasts. The cells were subjected to concentrations of 50µM and 500µM of 8Br-cAMP or db-cAMP, following the methodology previously outlined ^26^. These analogs activate the canonical PKA and EPAC pathways. After 48 hours of exposure in a serum-free medium, we observed a dose-dependent increase in ciliary abundance and cilia length across all cell lines. Notably, this increase in the average cilia length reached statistical significance at highest doses of 8Br-cAMP in the E36 cell line, which initially displayed the most severe phenotype at basal levels compared to I26 and control cells (2-way ANOVA with Tukey’s multiple comparison; n=2; p<0.05) (**Figure 6A and B; Figure S5**).

**Figure 6:**
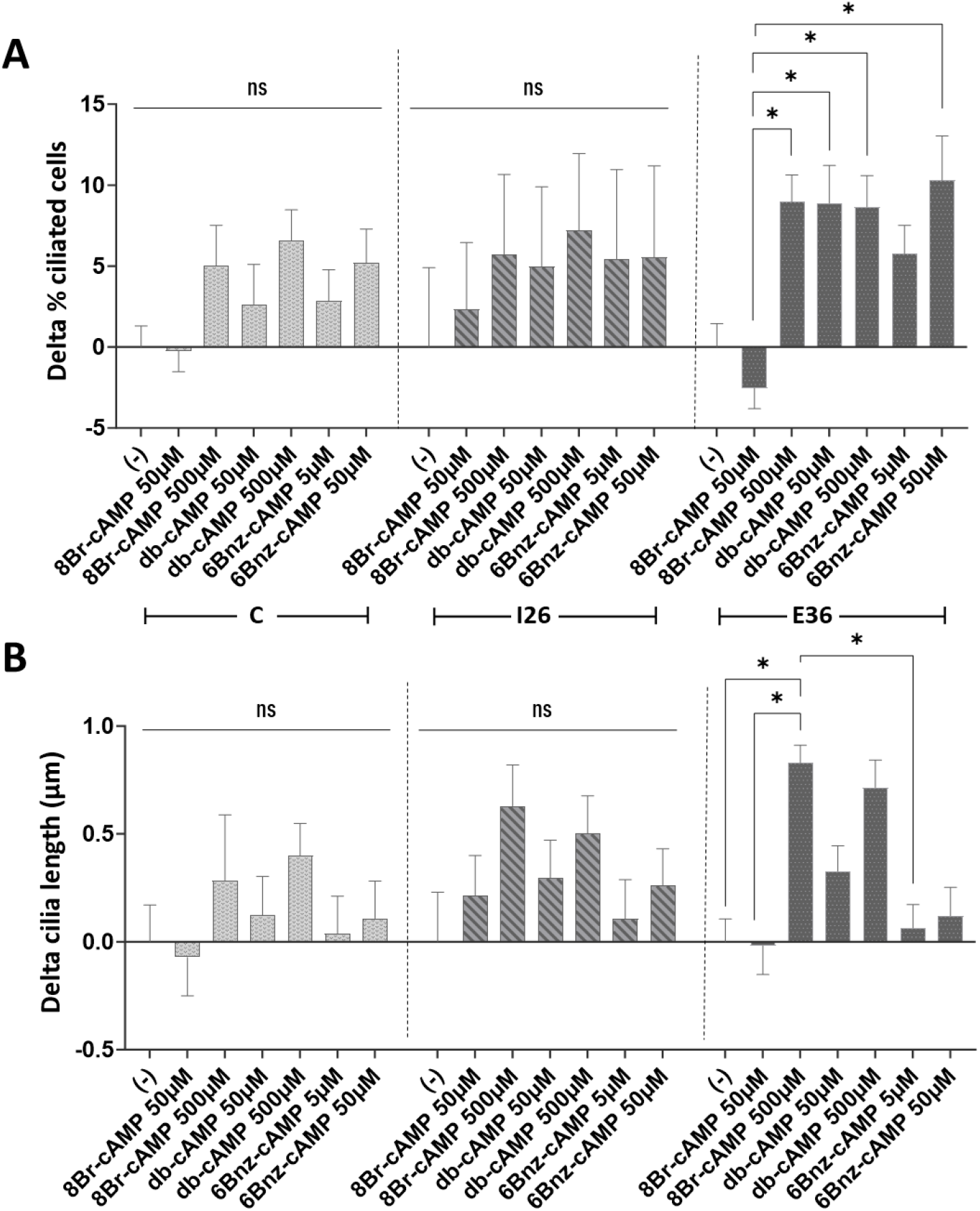
Impact of cAMP analogues on the ciliary phenotype of control and mutant fibroblasts. Serum-starved control (C: pool of C1 and C3) and mutant (I26, E36) fibroblasts were subjected to a high-content ciliogenesis assay. The cells were exposed to serum-free medium only (-) for an additional 48 hours or treated for 48 hours with cAMP analogs: 8-Bromo-cAMP (8Br-cAMP) at concentrations of 50 µM and 500 µM, dibutyryl-cAMP (db-cAMP) at concentrations of 50 µM and 500 µM, and 6-benzene-cAMP (6Bnz-cAMP) at concentrations of 5 µM and 50 µM, all diluted in water. The percentage of ciliated cells (**A**) and average cilia length (**B**) were automatically determined. The values represent the difference between each drug condition and the basal condition (serum-free medium only), as measured by automatic analysis. The bars indicate the mean ± SEM from 2 independent replicates. An ordinary two-way ANOVA was performed with a Tukey’s multiple comparison test within each cell line. P-values <0.05 is denoted by *; ns or no indication = not significant.

In addition, the cells were exposed to 5µM and 50µM of 6Bnz-cAMP, as indicated previously ^27^. This compound is specifically designed to stimulate the PKA pathway ^28,29^. We observed a dose-dependent trend towards increased ciliary abundance and cilia length across all cell lines, without reaching however statistical significance at the study doses compared to the basal condition (**Figure 6A and B; Figure S5**).

### Taprenenpag enhances photoreceptor outer segment formation and light response in a *Cep290* mouse model of retinal degeneration

To evaluate the *in vivo* impact of Taprenepag, we employed a mouse model with a homozygous in-frame deletion of exon 35 (equivalent to exon 36 in humans; *Cep290^del^*^36^*^/del^*^36^). This mouse model displays a normally developed retina but with rudimentary outer segments (OS) at eye opening (14 post-natal days, P14), leading to significantly altered ERG responses to both dim and bright light. By one month of age (P30), the outer nuclear layer (ONL) of the retina, which contains photoreceptor nuclei, is reduced by half, and by the age of 4 months, almost all nuclei are lost (Barny et al., In Preparation).

Before treatment, we assessed the retinal exposure following a single intraperitoneal injection of 8mg/kg Taprenepag in 2-month-old wild-type mice. We measured the concentrations of Taprenepag in plasma, retina, and vitreous humor at various timepoints (0.5, 1, 2, 4, 8, 24, 72, and 168 hours post-injection, pi). Taprenepag levels in plasma peaked at 1600ng/µL at 1 hour pi and remained substantial from 0.5 to 4 hours pi (**Figure 7A**). Regarding ocular tissues, Taprenepag was swiftly detected in the vitreous humor and retina within the first hour after injection, with concentrations of 0.016 ng/tissue and 0.057 ng/tissue at 0.5 hours post-injection, respectively (**Figure 7B**). This underscores the rapid delivery of the compound to the targeted tissues through intraperitoneal administration in less than an hour.

**Figure 7:**
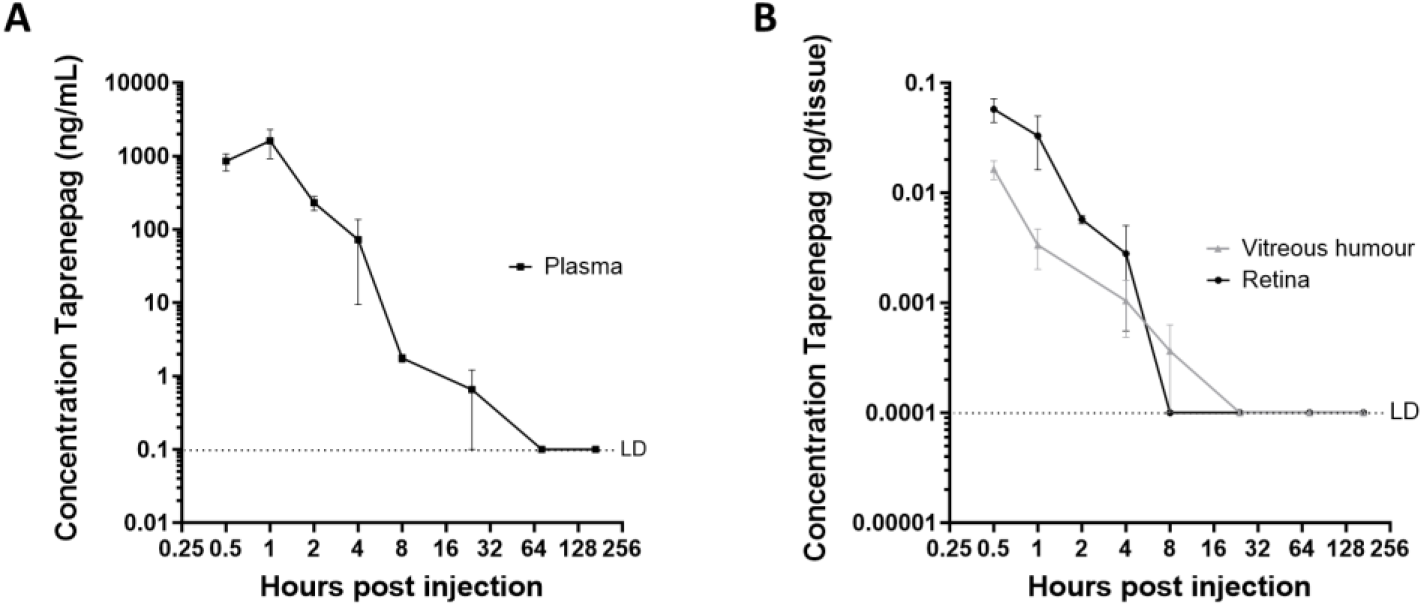
Measurement of Taprenepag levels in plasma and ocular tissues following a single intraperitoneal injection of 8mg/kg Taprenepag in WT Mice. Taprenepag concentration (ng/mL) over time post-injection in **A)** the plasma and **B)** the retina (dark) and vitreous humor (light grey) of WT mice. Bars represent the mean ± SEM from 4 mice. LD denotes the limit of detection.

Intraperitoneal injections of Taprenepag at a dosage of 18 mg/kg or vehicle (Solutol HS15) were administered every three days from postnatal day 6 (P6) to 27. We then evaluated the effects on outer segment (OS) formation, opsins content and ERG responses as well as cell loss at the age of P30.

Through histological analysis, we observed a reduction of the relative thickness of the outer nuclear layer (ONL) in both treated and untreated mutant animals in comparison with wildtype mouse. However, the reduction was a little less pronounced in the treated group as compared to the untreated group (Mann-Whitney U-test; n ≥ 5 mice; p < 0.05), suggesting a potential neuroprotective effect of the treatment. More interestingly, we observed that the average relative thickness of the layer composed of the inner and outer segments (IS/OS) was markedly increased (Mann-Whitney U-test; n ≥ 5 mice; p < 0.05) in mice exposed to Taprenepag as compared to the vehicle, strongly supporting improved OS formation (**Figure 8A and B**). Furthermore, through immunostaining of the rod opsin (rhodopsin), we observed a greater abundance of photopigments in the IS/OS of mice exposed to Taprenepag compared to the vehicle (Mann-Whitney U-test; n ≥ 3 mice; p<0.001) (**Figure 8C and D**). In line with the elevated levels of rhodopsin in the outer segments (OS), we observed higher amplitudes of the scotopic ERG a-waves, which measure the rod specific-response to light in mice exposed to Taprenepag compared to those exposed to the vehicle. This difference was particularly notable at the highest light intensities (**Figure 8E**). This improvement in the rod response, is statistically evident by the physiological amplification of the electrophysiological response carried out by the bipolar cells, which integrate the responses of the photoreceptors and are measured by the b-wave (2-way ANOVA with Sidak multiple comparison; n = 6 mice; p<0.01 and p<0.05 at 1 and 3 candela.s/m^2^, respectively) (**Figure 8E**).

**Figure 8:**
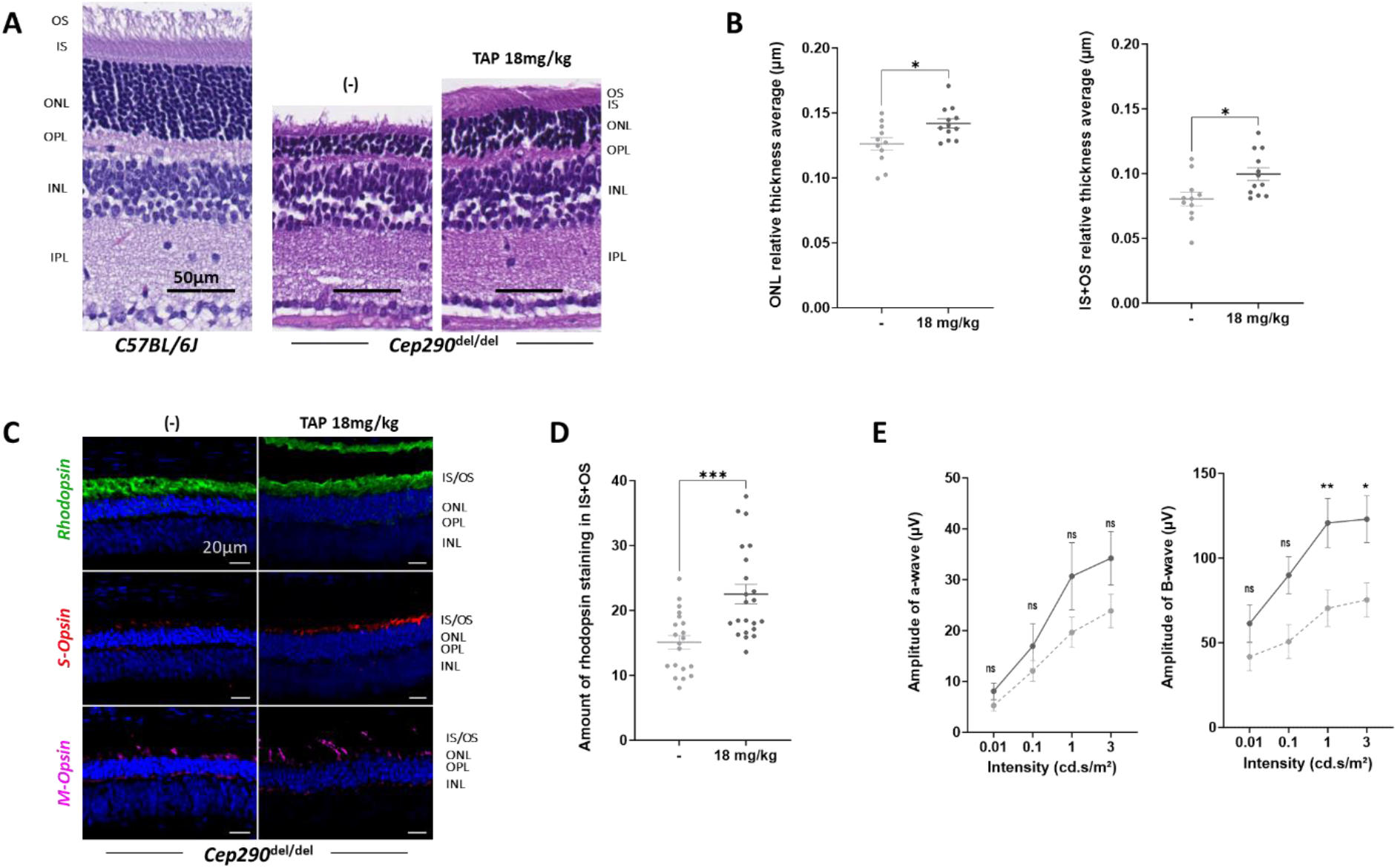
Impact of repeated intraperitoneal injections of Taprenepag on the structure and function of *Cep290^del36/del36^* mouse retina. **A.** Representative images of H&E retinal staining from P30 non-treated C57BL/6J and *Cep290^del^*^36^*^/del^*^36^ mice treated with vehicle (-) or 18 mg/kg Taprenepag (TAP 18mg/kg). Scale bar = 50µm. OS, outer segment; IS, inner segment; ONL, outer nuclear layer; OPL, outer plexiform layer; INL, inner nuclear layer; IPL, inner plexiform layer. **B.** Quantification of the Outer Nuclear Layer (ONL) & Inner Segment (IS) + Outer Segment (OS) thickness of the retina from P30 *Cep290^del^*^36^*^/del^*^36^ mice treated with vehicle (light grey) or 18 mg/kg Taprenepag (dark grey). Each point refers to the average of the layer thickness normalized by the total thickness of the retina for one eye. Bars correspond to the mean ± SEM from ≥ 5 animals. A Mann-Whitney U-test was performed, * indicates a p-value <0.05. **C.** Representative images of P30 *Cep290^del^*^36^*^/del^*^36^ mice retina following treatment with vehicle (-) or 18 mg/kg Taprenepag (TAP 18mg/kg). Rhodopsin labeling in green, S-opsin in red, and M-opsin in purple. Nuclei are stained with DAPI (Blue). Scale bar = 20µm. **D.** Quantification of the amount of Rhodopsin in the IS+OS from P30 *Cep290^del^*^36^*^/del^*^36^ mice treated with vehicle (light grey) or 18 mg/kg Taprenepag (dark grey). Each point refers to one field from one eye. Bars correspond to the mean ± SEM of 6 fields from ≥ 3 mice. A Mann-Whitney U-test was performed, *** indicates a p-value <0.001. **E.** Dark-adapted (Scotopic) ERG recordings in P30 *Cep290^del^*^36^*^/del^*^36^ mice after repeated intraperitoneal injections of 18mg/kg Taprenepag (dark grey) or vehicle (dotted light grey). Amplitude of the a-wave (Left) and B-wave (Right). Bars correspond to the mean ± SEM from 6 mice (left and right eyes). An ordinary two-way ANOVA was performed with Sidak’s multiple comparison test. P-values <0.01 and <0.05 are respectively indicated by ** and *, ns means no significance.

To evaluate the impact of Taprenepag on cone photoreceptors, which make up only 1-3% of all photoreceptors in the mouse retina and require high-intensity light stimuli to function, we conducted immunostaining for the S and M opsins and ERG recordings upon bright light stimulation. We observed enhanced immune S and M opsins signals in mice exposed to Taprenepag in comparison to the vehicle (**Figure 8C**). Additionally, we detected a trend towards an increase in the photopic b-wave, which measures the contribution of cones to bipolar cells’ activity (**Figure S6**).

Taken together, these findings indicate that the intraperitoneal administration of Taprenepag mitigates retinal degeneration in *Cep290^del^*^36^*^/del^*^36^ mice by enhancing outer segment (OS) formation and increasing the availability of light-sensitive photopigments. This results in a subtle yet significant enhancement in retinal function, noticeable through the recording of the magnified responses from secondary retinal neurons (bipolar cells) elicited by the highest-light-sensitive photoreceptors (rods) when exposed to intense light.

## DISCUSSION

The research discussed here sheds light on a promising approach to address ciliary defects associated with *CEP290* mutations, which present significant challenges in inherited retinal diseases.

The utilization of skin-derived fibroblasts as a surrogate model system for studying the cellular and molecular mechanisms underlying ciliary dysfunctions and screening therapeutic compounds is gaining prominence, especially in the context of retinal anomalies. One of the key advantages of using skin-derived fibroblasts is their accessibility through a relatively minimally invasive procedure. Moreover, the use of patient-derived fibroblasts allows for personalized investigations, as each patient may possess unique genotypes, even when they share the same mutations.

Our study introduced cell lines containing two prevalent *CEP290* mutations: the deep intronic c.2991+1655A>G variant and the nonsense c.4723A>T mutation, responsible for approximately 20% and 2.5% of non-syndromic Leber congenital amaurosis (LCA) cases in children, respectively ^17,30^. Notably, the c.2991+1655A>G mutation has previously shown the ability to express up to 50% of the normal *CEP290* transcript, while the c.4723A>T mutation sporadically leads to exon skipping, resulting in a slightly shortened but still functional transcript ^1,17^. Additionally, we included fibroblasts from a fetus affected by Meckel-Gruber syndrome (MKS), harboring compound heterozygous mutations in *CEP290*, which are anticipated to result in the absence of detectable CEP290 protein ^31^. This comprehensive selection of mutations allowed us to observe a connection between *CEP290* genotypes and the degree of cilia abundance reduction. Moreover, it unveiled a spectrum of effects on ciliary lengths, ranging from highly variable to even opposing outcomes. Importantly, our study demonstrated that exposure to Taprenepag improved these features across all cell lines, including both the control and mutant groups. This observation not only underscores the effectiveness of Taprenepag, irrespective of the *CEP290* genotype, but also suggests that its effects are independent of CEP290, potentially making this a mutation-agnostic treatment.

The precise mechanism through which Taprenepag enhances cilia formation in *CEP290* mutants remains enigmatic. Prostaglandin functional analogs like Taprenepag, exert their effects by activating specific prostaglandin receptors, resulting in increased cAMP levels. cAMP plays a crucial role as a second messenger in the regulation of cilia biogenesis and elongation. It exerts its influence on various cellular processes, including protein phosphorylation through the activation of protein kinase A (PKA), which has been reported to enhance anterograde ciliary trafficking ^32^. Additionally, cAMP is involved in the modulation of microtubule dynamics, regulation of gene expression, and interactions with other signaling pathways. The role of cAMP may vary depending on cell type, subcellular localization and context. Several studies have suggested that the primary cilium forms an independent cAMP sub-compartment, supported by the unique composition of the ciliary membrane housing various receptors, including rhodopsin-type G protein-coupled receptors (GPCRs), and signaling pathways that cross paths with cAMP signaling ^33^.

Taprenepag is a potent and highly selective non-prostanoid agonist, demonstrating a significant preference for the EP_2_ receptor over the EP_1_, EP_3_, and EP_4_ receptors ^22^. In this study, we have confirmed the expression of *PTGER_2_* in human fibroblasts and observed a decrease in its expression following exposure to Taprenepag. This observation, consistent with the downregulation of *PTGER_2_* expression upon endogenous PGE_2_ binding in human monocytic cells ^34^, strongly substantiates the interaction between Taprenepag and the EP_2_ receptor in human fibroblasts.

Our findings demonstrate that exposure to Taprenepag significantly enhances both total cAMP synthesis and the formation and elongation of cilia. While the classification of EP_2_ as a ciliary GPCR has not been established, the mechanism by which Taprenepag influences ciliation, whether through its impact on ciliary or cellular cAMP synthesis, warrants further investigation. Notably, Taprenepag exhibited the ability to promote cilia formation in E14/E32 fibroblasts, the vast majority of which (> 90%) lacked a primary cilium (**Figure 1**). CEP290 is a key component of the NPHP/MKS ciliary module located at the ciliary transition zone (TZ) ^35^. Here, it serves as a critical element in maintaining the structural integrity and barrier function of the cilia base. This function controls the flow of proteins and molecules into and out of the ciliary compartment, thereby governing ciliary signaling and maintaining overall ciliary structure. In the absence of CEP290, the basal body assembly remains intact; however, there is a dramatic impairment in axonemal elongation. This raises questions about the primary role of ciliary cAMP signaling in the earliest stages of cilia formation. It is reasonable to hypothesize that initially, the intracellular pool of cAMP contributes to cilia formation by influencing cAMP-dependent cellular signaling pathways. Subsequently, the ciliary cAMP pool likely plays a key role in regulating axonemal elongation. Further supporting a role of intracellular cAMP is the enhanced cilia formation upon exposure to cAMP analogs in the culture medium. It is noteworthy that despite expression of *PTGER_4_* mRNA in human fibroblasts, the involvement of the EP_4_ receptor, known as a ciliary GPCR ^32^, in the response to Taprenepag appears unlikely.

This conclusion is strengthened by the remarkable selectivity of Taprenepag for EP_2_. Furthermore, the absence of a synergistic effect observed upon exposure to Alprostadil, which exhibits similar affinities for both EP_2_ and EP_4_ receptors, reinforces this conclusion.

One of our key findings is the effectiveness of Taprenepag in increasing intracellular cAMP levels, even at significantly lower doses than Forskolin. This raises intriguing questions about the mechanisms behind Taprenepag’s potency. Importantly, both compounds exhibit a dose-dependent increase in cAMP concentration, ruling out the possibility of EP_2_ or adenylate cyclase (AC) saturation by their ligands. The greater potency of Taprenepag may stem from several factors, including enhanced cell permeability and cellular availability, increased binding or activation of its target EP2 receptors in comparison to Forskolin’s interaction with AC, and/or the potential involvement of an upstream amplification mechanism within the cAMP pathway. Exploring these aspects could lead to the development of more targeted and potent therapies for addressing ciliary defects associated with *CEP290* mutations, presenting an exciting avenue for further research.

While skin-derived fibroblasts are a valuable tool for initial screening and mechanistic studies, translating findings to the complex *in vivo* environment of the retina can be challenging. Cellular responses in fibroblasts may not fully recapitulate the conditions in retinal cells, necessitating further validation in relevant models.

In this study, we demonstrate that repetitive intraperitoneal injections of Taprenepag in *Cep290^del^*^36^*^/del^*^36^ mice with retinal degeneration yield a promising neuroprotective effect on the outer nuclear layer (ONL). Notably, this parallels a previous study where intraperitoneal administration of Taprenepag mitigated retinal degeneration in *Nphp1*^-/-^ mice ^36^. Our study extends these findings by revealing enhanced OS and an increase in opsin content, which aligns closely with our observations in fibroblasts, indicating improved cilia formation. This enhancement likely results from EP_2_ stimulation by Taprenepag, as supported by the presence of EP_2_ expression in various neuronal cells in both mouse and human retinas, including photoreceptors ^37–39^.

Enhanced outer segment formation may mitigate the accumulation of toxic proteins in the inner segments (IS), including opsins and other visual signaling components, which have been associated with retinal degeneration in various mouse models. However, multiple mechanisms may contribute to these positive effects. Notably, EP_2_ -cAMP pathway activation in Müller glial cells could lead to the release of neurotrophic factors, promoting neuroprotection ^39^, or it might enhance retinal vascularization through vasodilation in retinal vascular smooth muscles, potentially delaying photoreceptor degeneration ^40^.

Extending these findings to clinical applications holds promise, particularly when considering the measurable impact of the treatment on electroretinogram (ERG) results. It is important to note that the detection threshold for light responses is elevated in certain cases, as demonstrated by the absence of or recordable ERG in individuals with low but still measurable visual acuity, such as children carrying mutations in genes like *RPE65* ^41^.

Furthermore, the slower rate of photoreceptor cell loss in LCA10 patients with *CEP290* mutations compared to the *Cep290^del^*^36^*^/del3^*^6^ mouse model is noteworthy. Specifically, dormant cones have been detected in the central retina of *CEP290* patients up to the age of 30 years ^13^, which roughly corresponds to 3-9 months of age in mice. Consequently, the observed neuroprotective and functional benefits in *Cep290^del^*^36^*^/del3^*^6^ mice suggest the potential to extend the therapeutic window and enhance visual function for LCA10 patients through the use of Taprenepag.

In summary, our study offers promising insights into addressing ciliary defects associated with *CEP290* mutations, a challenging aspect of inherited retinal diseases. Leveraging skin-derived fibroblasts as a practical model system, we demonstrated the effectiveness of Taprenepag in improving ciliogenesis across various *CEP290* mutations, suggesting its potential as a “mutation-agnostic” therapy. While the exact mechanism remains unclear, cAMP signaling likely plays a key role, impacting cilia biogenesis and reducing toxic protein accumulation in photoreceptors. Moreover, our findings extend to clinical applications, where Taprenepag shows measurable impacts on electroretinogram results in the mouse, presenting a potential opportunity to enhance visual function for LCA10 patients with *CEP290* mutations. This research lays a foundation for the development of targeted therapies for ciliary defects linked to *CEP290* mutations, addressing a critical need in the field of inherited retinal diseases.

## MATERIAL AND METHODS

### Fibroblasts

Primary fibroblasts used in this study include lines carrying biallelic *CEP290* mutations of increasing severity (I26, E36, E14/E32 lines, respectively) and control lines (C1, C2, C3).

### Cell culture

Primary fibroblasts were isolated through selective trypsinization and cultivated at 37°C with 5% CO2 in “complete medium,” which consists of Opti-MEM Glutamax I medium (Invitrogen, Carlsbad, CA, USA), supplemented with 10% fetal bovine serum (Invitrogen), 1% Ultroser-G serum substitute (Sartorius, Aubagne, France), and 1% penicillin/streptomycin (Invitrogen). The cells were cultured in 75 cm^2^ flasks and used at less than 15 passages.

### RNA and protein analysis

Following 24 hours in serum-free medium, cells were exposed to either 0.2µM of Taprenepag (Edelris, Lyon, France), in DMSO, or DMSO alone for an additional 24 hours before being harvested. Total mRNA extraction was conducted using the RNeasy Micro kit (Qiagen, Les Ulis, France) in accordance with the manufacturer’s guidelines. The concentration and purity of the total mRNA were determined using a Nanodrop-2000 spectrophotometer (ThermoScientific). Subsequently, the RNAs were reverse transcribed into cDNA using the Verso cDNA Kit (ThermoFisher Scientific, Asnières-sur-Seine, France) and amplified by PCR using the GoTaq Master Mix (Promega, Charbonnières les Bains, France). We employed specific primers for *PTGER_1_*, *PTGER_2_*, *PTGER_3_*, and *PTGER_4_* (**Supplementary Table 1**).

For real-time quantitative PCR (RT-qPCR), cDNA amplification was performed following a previously described protocol ^17^ using the same primers as for PCR. To normalize the data, human β-glucuronidase (GUSB, NM_000181.3) and P0 large ribosomal protein (RPLP0, NM_001002.3) mRNAs were utilized, and negative control reactions (no template, NTC) were included. Data analysis followed established procedures ^17,24^.

Protein extraction and Western Blot analysis were performed using previously established procedures ^17,24^.

### Intracellular cAMP Quantification

To measure the drug-induced increase in intracellular cAMP, we employed the cAMP-Glo Assay (Promega). Primary fibroblasts were initially seeded in a 96-well plate at a density of 100,000 cells/cm^2^ (32,000 cells/well) in complete medium. The following day, cells underwent serum starvation by replacing the medium with Opti-MEM Glutamax I medium. After 24 hours, cells were pre-incubated for 10 minutes at 37°C in an “incubation buffer,” consisting of Opti-MEM Glutamax I medium supplemented with 500µM isobutyl-1-methylxanthine (IBMX, Sigma Aldrich, Saint Quentin Fallavier, France) and 100µM 4-(3-butoxy-4-methoxy-benzyl) imidazolidone (Ro20-1724, Sigma Aldrich). IBMX and imidazolidone act as phosphodiesterase inhibitors, preventing cAMP hydrolysis.

Following the pre-incubation step, cells were exposed to either DMSO or the drug of interest, which was diluted to the desired concentration in DMSO, within the “incubation buffer” for another 10 minutes at 37°C. A cAMP standard curve was generated through successive dilutions, encompassing 12 concentrations ranging from 0 to 4µM. Each condition, including the cAMP standard curve, was evaluated in triplicate. Subsequent assay steps were carried out following the manufacturer’s instructions, and luminescence measurements were recorded using the Victor luminometer (Perkin Elmer, Villebon sur Yvette, France).

### Ciliogenesis assay

In this ciliogenesis assay, we seeded 150,000 cells per well on glass coverslips in a 12-well plate. After 24 hours of incubation in complete medium, ciliogenesis was induced by switching to serum-free Opti-MEM Glutamax I medium. To assess basal ciliogenesis, cells were starved for 48 hours before fixation. For drug experiments, cells were serum-starved for 24 hours and then incubated for an additional 24 hours with drugs diluted in Opti-MEM Glutamax I. Taprenepag (Edelris) and Forskolin (Sigma Aldrich) were initially diluted in DMSO to achieve a final DMSO concentration of 0.04% in the well. After fixation with cold methanol (−20°C for 7 minutes), cells were blocked for 1 hour at room temperature using PBS 1X buffer supplemented with 5% Normal Goat Serum (Sigma Aldrich), 3% Bovine Serum Albumin (BSA, Sigma Aldrich), and 0.5% Triton X-100. The ciliary membrane was stained with rabbit anti-ARL13B antibody (1:1000, Proteintech, Manchester, UK), and the basal body was stained with mouse anti-gamma tubulin antibody (clone GTU-88 1:5000, Sigma Aldrich). Following overnight incubation at 4°C with primary antibodies diluted in PBS 1X buffer supplemented with 3% BSA and 0.1% Triton X-100, cells were incubated for 1 hour at room temperature with secondary antibodies (Alexa555 - donkey anti-rabbit and Alexa 488 - donkey anti-mouse, 1:2000, Thermofisher Scientific) and DAPI diluted in the same buffer. Coverslips were mounted on slides using Fluoromount medium (Sigma Aldrich), and immunofluorescence images were acquired using a Zeiss LSM700 confocal microscope. Exposure time and settings were kept consistent for all samples to enable sample comparison. Five fields were captured for each sample using the 20X objective, and the number of Z-stacks collected varied among samples, optimized for maximum fluorescent signal capture. Images were then combined into a single picture using the Fiji software Z-project tool with the maximum intensity setting ^42^. Finally, the ciliary phenotype, including abundance and average cilia length, was manually analyzed in each field using the counting and measuring tools of Fiji.

### Ciliogenesis assay in high content condition

In the high-content immunocytochemistry ciliogenesis assay, cells were initially seeded in 96-well plates (Cell Carrier Ultra Plates, Perkin Elmer) at a density of 100,000 cells/cm^2^. Following a 24-hour incubation in complete medium, ciliogenesis was induced through serum starvation and exposure to Opti-MEM Glutamax I Medium during 24 hours. Subsequently, cells underwent various drug experiments: for cAMP analogs (8Br-cAMP, db-cAMP, and 6Bnz-cAMP, Tocris Bioscience, Bristol, UK), they were incubated for 48 hours with the appropriate concentrations (50 and 500µM for 8Br-cAMP and db-cAMP, or 5 and 50µM for 6Bnz-cAMP); for PGE2 agonists (Taprenepag and Alprostadil, Edelris), they were incubated for 24 hours with a final DMSO concentration of 0.04% and a compound concentration of 0.2µM. Fixation was performed for 7 minutes using cold methanol (−20°C). After fixation, cells were blocked for 1 hour at room temperature, either with PBS 1X buffer supplemented with 10% Normal Goat Serum (NGS) and 0.1% Tween 20 or with 5% NGS, 3% Bovine Serum Albumine (BSA), and 0.5% Triton X-100. Staining was carried out using a rabbit anti-ARL13B antibody (1:1000, Proteintech) for the ciliary membrane and a mouse anti-gamma tubulin antibody (clone GTU-88 1:5000, Sigma Aldrich) for the basal body. Primary antibody incubation was performed overnight at 4°C, followed by secondary antibody incubation (Alexa555 - Goat anti Rabbit and Alexa 488 - Goat anti Mouse, 1:2000, Thermofisher Scientific) and DAPI staining at room temperature for 1 hour in the same buffer. Imaging was performed using the Perkin Elmer Opera high-throughput imager. The analysis included acquiring images using a 5X objective to detect DAPI and assess cell seeding, and a 40X water objective lens to capture images of cilia using DAPI, Alexa 488, and Alexa 555. Images were processed using Harmony 4.9 Perkin Elmer software, and nuclei, basal bodies, and cilia of at least 150 cells were quantified using image segmentation.

### Mice breeding and intraperitoneal injections

The mice were bred and maintained at the LEAT Facility of Imagine Institute under a 12-hour light/dark cycle.

Intraperitoneal injections of 18mg/kg of Taprenepag, diluted in either Solutol HS15 5% or Solutol HS15 5% alone (Drugabilis, Villejust, France), were administered every 3 days to male and female mice carrying a *Cep290* gene deleted of exon 35 (36 in human) in homozygosity (*Cep290^del^*^36^*^/del^*^36^) from P6 to P27. Each treatment group included 6 *Cep290^del^*^36^*^/del36^*mice. At P30, electroretinograms were recorded from all *Cep290^del^*^36^*^/del36^*mice before euthanasia, which was performed by cervical dislocation, followed by the collection of tissue samples.

### Electroretinograms

Electroretinograms (ERGs) were recorded using a commercial diagnostic system (Celeris; Diagnosys LLC, Lowell, USA). Mice were subjected to overnight dark adaptation and then anesthetized through intramuscular injection of ketamine (120 mg/kg) and xylazine (16 mg/kg). Pupils were dilated with 0.5% tropicamide and 10% phenylephrine drops, followed by the application of sterile ophthalmic gel on the corneal surface to ensure proper electrical contact and maintain corneal integrity. Throughout the ERG procedure, animals were kept on a warming support within the Celeris apparatus to maintain their body temperature at 38°C. Stimuli and recordings were generated using Celeris electrode stimulators, with a subcutaneously inserted ground electrode. The dark-adapted ERG protocol involved four steps with increasing stimulus strengths, ranging from 0.01 to 3 cd.s/m^2^. The light-adapted ERG protocol was composed of two steps with light intensities of 3 and 10 cd.s/m^2^.

### Retina Histology

Freshly enucleated eyes were fixed in a phosphate-buffered saline (PBS) solution containing 4% paraformaldehyde (PFA, Santa Cruz, Athis-Mons, France) at 4°C for 12 hours. Subsequently, at the Imagine Institute’s histology platform, the eyes underwent rehydration using a series of ethanol gradients with the assistance of an automated tissue processor (ASP300S, Leica). Afterward, they were embedded in paraffin and longitudinally sectioned using microtomes (2 x HM 340E, Microm France) to produce serial sections encompassing the entire retina.

Each section underwent staining with haematoxylin and eosin before being scanned utilizing a commercial imaging system (NanoZoomer S210, Hamamatsu). Analysis was carried out using NDPview software, during which the thickness of the retina’s outer nuclear layer and inner and outer segments layer was measured at various points along the retina (at distances of 0.25, 0.5, 0.75, 1, 1.25, 1.5, 1.75, 2, and 2.25mm from the optic nerve) and normalized to the thickness of the total retina. The average relative thickness for each layer was then calculated for each eye.

### Immunohistostaining

Paraffin-embedded sections were subjected to deparaffinization and rehydration through Histoclear and descending ethanol gradients. Antigen retrieval was performed by incubating the sections in a solution containing 10mM Tris-HCl (Tris(hydroxymethyl) amino-methane) at pH 9.2, 2mM EDTA, and 0.01% Tween for 7 minutes at 900W in a microwave. Subsequently, the sections were blocked for 1 hour using a blocking solution comprising 0.1% Tween, 50mM NH4Cl, and 1% BSA in PBS 1X.

Primary antibodies were prepared in the blocking solution, and the slides were incubated overnight at 4°C in a humidifying chamber. The antibodies used included anti-S Opsin rabbit antibody (1:100, Invitrogen), anti-M Opsin rabbit antibody (1:1000, Invitrogen), and anti-Rhodopsin mouse antibody (1:500, Novus Biologicals, Noyal Châtillon sur Seiche, France). Following primary antibody incubation, the sections were washed three times with PBS 1X and then incubated for 1 hour with donkey anti-mouse (or anti-rabbit) Alexa Fluor 488nm (or 555nm) secondary antibodies (1:200, Thermofisher Scientific). DAPI (Roche, Boulogne-Billancourt, France) was diluted with PBS 1X to a final concentration of 1.25µg/mL and used to label the nuclei of the sections.

After additional PBS washing, the sections were mounted with Fluoromount medium (Sigma Aldrich) under glass coverslips and visualized using a Zeiss LSM700 confocal microscope. For each eye, three fields on each side of the optic nerve were captured with the 20X objective. The number of Z-stacks collected varied between samples but was optimized to capture maximum fluorescent signals.

Initial image preprocessing involved applying a sum intensity Z projection using a Fiji v2.14.0/1.54f macro ^42^. Quantitative analysis was then conducted using QuPath v0.4.3 software ^43^. The outer nuclear layer (ONL) and inner and outer segments layer (IS+OS) on preprocessed images were manually segmented and annotated in each field based on DAPI and rhodopsin staining. To distinguish positive rhodopsin signal from negative, a machine learning pixel classification using the random forest algorithm was applied to various fields extracted from representative images. The model created was then used for annotations on all images. The amount of rhodopsin in the IS+OS was determined as the total area of positive pixels detected by the model in the IS+OS layer, divided by the length of the analyzed section.

### Pharmacokinetics study

Taprenepag was obtained from MedChem Express (Batch#20534, Sollentuna, Sweden), and injected formulated in Solutol HS15 5% in buffer pH7.4 in CB57BL6/J males of approximately 8 weeks of age. Animals were selected on good health and homogenous body weights. Timepoints after intraperitoneal injection were 0,5h (-/+3min), 1h(-/+6min), 2h (-/+12min), 4h (-/+24min), 8h(-/+50min), 24h(-/+2h), 72h(-/+7h), 168h(-/+16h), 4 mice per timepoint. After euthanasia, pooled retina and vitreous were dissected from both eyes and weighted, snap-frozen and stored in Eppendorf’s ™ protein low binding 1.5mL microtubes, and stored at −80°C +/− 15°C. Blood of each animal was centrifuged at 2000g, 10min at 4°C. Approximately 75µl of plasma were sampled and stored at −80°C +/− 15°C. All animal procedures were done at Iris Pharma (la Gaude, France); all standard opening procedures and protocols described were reviewed by Iris Pharma Internal Ethics Committee (DPA66). All animals were treated according to the Directive 2010/63/UE European Convention for the Protection of vertebrate Animals used for Experimental and Other Scientific Purposes. Plasma concentrations were determined using protein precipitation for sample preparation followed by Liquid chromatography/mass spectrometry (LC/MS) detection. Retinas were prepared by acid maceration. Retina and vitreous humor concentrations were determined using liquid-liquid extraction for sample preparation followed by LC-MS detection, using a triple quadruple API6500 (Thermofisher Scientific).

### Figures

The figures were generated using the FigureJ plugin in Fiji. ^44^

### Statistics

Statistical analyses were conducted using GraphPad Prism version 10.0.0 for Windows, a software developed by GraphPad Software in San Diego, California, USA, which is available at www.graphpad.com.

### Study approval

All animal procedures were conducted in accordance with ethical principles and received approval from the French Ministry of Research (APAFIS# 2023092513468026), in full compliance with the guidelines for animal experiments in France.

## AUTHOR CONTRIBUTIONS

JMR and JPA conceived and directed the project. FM and IB were responsible for research execution, data acquisition and interpretation. LBR contributed to data acquisition and interpretation. SM, IP and EC contributed to the experiments. LFT contributed to murine models creation. NG designed the methodology of rhodopsin quantification. JK and TAB provided mutant cells. Manuscript preparation was done by FM, with revisions provided by IB, JPA, LBR and JMR.

## ACKNOWLEDGMENTS

We are grateful to the Association Retina France (Grant to IP and JMR), the ANRT (Association Nationale Recherche Technologie, Grant to FM), the SGPI (General Secretary for Investment in France) with the program RHU (Research Hospital University C’IL LICO vague 3), the BPI (Bank For Innovation) for the supporting award I-Lab 2022, and InsermTransfert for the HHSF (Human Health Start Up Factory program). We gratefully acknowledge the patients that donated skin biopsies for this study. We gratefully acknowledge members of the Histology and LEAT platforms of Imagine institute for excellent assistance with histology and mouse care.

## REFERENCES

1. Gerard X, Perrault I, Hanein S, et al. AON-mediated exon skipping restores ciliation in fibroblasts harboring the common leber congenital amaurosis CEP290 mutation. Mol Ther - Nucleic Acids. 2012;1(6):e29. doi:10.1038/mtna.2012.21

2. Stephen P. Daiger, Lori S. Sullivan SJB. RetNet, the Retinal Information Network. Investigative Ophthalmology & Visual Science. Published 1998. Accessed April 10, 2023. https://web.sph.uth.edu/RetNet/

3. Sung CH, Chuang JZ. The cell biology of vision. J Cell Biol. 2010;190(6):953–963. doi:10.1083/jcb.201006020

4. Pearring JN, Salinas RY, Baker SA, Arshavsky VY. Protein sorting, targeting and trafficking in photoreceptor cells. Prog Retin Eye Res. 2013;36:24–51. doi:10.1016/j.preteyeres.2013.03.002

5. Koenekoop RK. An overview of leber congenital amaurosis: A model to understand human retinal development. Surv Ophthalmol. 2004;49(4):379–398. doi:10.1016/j.survophthal.2004.04.003

6. Huang CH, Yang CM, Yang CH, Hou YC, Chen TC. Leber’s congenital amaurosis: Current concepts of genotype-phenotype correlations. Genes (Basel). 2021;12(8). doi:10.3390/genes12081261

7. Quinlan RJ, Tobin JL, Beales PL. *Chapter 5 Modeling Ciliopathies*. Primary Cilia in Development and Disease. Vol 84. 1st ed. Elesvier Inc.; 2008. doi:10.1016/S0070-2153(08)00605-4

8. Parfitt DA, Lane A, Ramsden CM, et al. Identification and Correction of Mechanisms Underlying Inherited Blindness in Human iPSC-Derived Optic Cups. Cell Stem Cell. 2016;18(6):769–781. doi:10.1016/j.stem.2016.03.021

9. Shimada H, Lu Q, Insinna-Kettenhofen C, et al. In Vitro Modeling Using Ciliopathy-Patient-Derived Cells Reveals Distinct Cilia Dysfunctions Caused by CEP290 Mutations. Cell Rep. 2017;20(2):384–396. doi:10.1016/j.celrep.2017.06.045

10. Boye SE, Huang WC, Roman AJ, et al. Natural history of cone disease in the murine model of Leber congenital amaurosis due to CEP290 mutation: Determining the timing and expectation of therapy. PLoS One. 2014;9(3). doi:10.1371/journal.pone.0092928

11. Chang B, Khanna H, Hawes N, et al. In-frame deletion in a novel centrosomal/ciliary protein CEP290/NPHP6 perturbs its interaction with RPGR and results in early-onset retinal degeneration in the rd16 mouse. Hum Mol Genet. 2006;15(11):1847–1857. doi:10.1093/hmg/ddl107

12. Cideciyan A V., Rachel RA, Aleman TS, et al. Cone photoreceptors are the main targets for gene therapy of NPHP5 (IQCB1) or NPHP6 (CEP290) blindness: Generation of an all-cone Nphp6 hypomorph mouse that mimics the human retinal ciliopathy. Hum Mol Genet. 2011;20(7):1411–1423. doi:10.1093/hmg/ddr022

13. Cideciyan A V., Jacobson SG. Leber congenital amaurosis (Lca): Potential for improvement of vision. Investig Ophthalmol Vis Sci. 2019;60(5):1680–1695. doi:10.1167/iovs.19-26672

14. Cideciyan A V., Aleman TS, Boye SL, et al. Human gene therapy for RPE65 isomerase deficiency activates the retinoid cycle of vision but with slow rod kinetics. Proc Natl Acad Sci U S A. 2008;105(39):15112–15117. doi:10.1073/pnas.0807027105

15. Cideciyan A V., Jacobson SG, Beltran WA, et al. Human retinal gene therapy for Leber congenital amaurosis shows advancing retinal degeneration despite enduring visual improvement. Proc Natl Acad Sci U S A. 2013;110(6):517–525. doi:10.1073/pnas.1218933110

16. Molinari E, Decker E, Mabillard H, et al. Human urine-derived renal epithelial cells provide insights into kidney-specific alternate splicing variants. Eur J Hum Genet. 2018;26(12):1791–1796. doi:10.1038/s41431-018-0212-5

17. Barny I, Perrault I, Michel C, et al. AON-mediated exon skipping to bypass protein truncation in retinal dystrophies due to the recurrent CEP290 c.4723A > T mutation. Fact or fiction? Genes (Basel). 2019;10(5). doi:10.3390/genes10050368

18. Cideciyan A V., Jacobson SG, Ho AC, et al. Durable vision improvement after a single intravitreal treatment with antisense oligonucleotide in CEP290-LCA: Replication in two eyes. Am J Ophthalmol Case Reports. 2023;32(June):101873. doi:10.1016/j.ajoc.2023.101873

19. Russell SR, Drack A V., Cideciyan A V., et al. Intravitreal antisense oligonucleotide sepofarsen in Leber congenital amaurosis type 10: a phase 1b/2 trial. Nat Med. 2022;28(5):1014–1021. doi:10.1038/s41591-022-01755-w

20. Burnight ER, Gupta M, Wiley LA, et al. Using CRISPR-Cas9 to Generate Gene-Corrected Autologous iPSCs for the Treatment of Inherited Retinal Degeneration. Mol Ther. 2017;25(9):1999–2013. doi:10.1016/j.ymthe.2017.05.015

21. Ruan GX, Barry E, Yu D, Lukason M, Cheng SH, Scaria A. CRISPR/Cas9-Mediated Genome Editing as a Therapeutic Approach for Leber Congenital Amaurosis 10. Mol Ther. 2017;25(2):331–341. doi:10.1016/j.ymthe.2016.12.006

22. Prasanna G, Carreiro S, Anderson S, et al. Effect of PF-04217329 a prodrug of a selective prostaglandin EP 2 agonist on intraocular pressure in preclinical models of glaucoma. Exp Eye Res. 2011;93(3):256–264. doi:10.1016/j.exer.2011.02.015

23. Molinari E, Ramsbottom SA, Srivastava S, et al. Targeted exon skipping rescues ciliary protein composition defects in Joubert syndrome patient fibroblasts. Sci Rep. 2019;9(1):1–13. doi:10.1038/s41598-019-47243-z

24. Barny I, Perrault I, Michel C, et al. Basal exon skipping and nonsense-associated altered splicing allows bypassing complete CEP290 loss-of-function in individuals with unusually mild retinal disease. Hum Mol Genet. 2018;27(15):2689–2702. doi:10.1093/hmg/ddy179

25. Tsuboi K, Sugimoto Y, Ichikawa A. Prostanoid receptor subtypes. Prostaglandins Other Lipid Mediat. 2002;68-69:535–556. doi:10.1016/S0090-6980(02)00054-0

26. Mori Sequeiros Garcia M, Gorostizaga A, Brion L, González-Calvar SI, Paz C. CAMP-activated Nr4a1 expression requires ERK activity and is modulated by MAPK phosphatase-1 in MA-10 Leydig cells. Mol Cell Endocrinol. 2015;408:45–52. doi:10.1016/j.mce.2015.01.041

27. Xiao LY, Kan WM. Cyclic AMP (cAMP) confers drug resistance against DNA damaging agents via PKAIA in CML cells. Eur J Pharmacol. 2017;794(November 2016):201–208. doi:10.1016/j.ejphar.2016.11.043

28. Christensen AE, Selheim F, De Rooij J, et al. cAMP analog mapping of Epac1 and cAMP kinase: Discriminating analogs demonstrate that Epac and cAMP kinase act synergistically to promote PC-12 cell neurite extension. J Biol Chem. 2003;278(37):35394–35402. doi:10.1074/jbc.M302179200

29. Leech CA, Chepurny OG, Holz GG. Epac2-Dependent Rap1 Activation and the Control of Islet Insulin Secretion by Glucagon-Like Peptide-1. Vol 84. 1st ed. Elsevier Inc.; 2010. doi:10.1016/B978-0-12-381517-0.00010-2

30. Den Hollander AI, Koenekoop RK, Yzer S, et al. Mutations in the CEP290 (NPHP6) gene are a frequent cause of leber congenital amaurosis. Am J Hum Genet. 2006;79(3):556–561. doi:10.1086/507318

31. Baala L, Romano S, Khaddour R, et al. The Meckel-Gruber syndrome gene, MKS3, is mutated in Joubert syndrome. Am J Hum Genet. 2007;80(1):186–194. doi:10.1086/510499

32. Jin D, Zhong TP. Prostaglandin signaling in ciliogenesis and development. J Cell Physiol. 2021;16(9):9–10. doi:10.1002/jcp.30659

33. Paolocci E, Zaccolo M. Compartmentalised cAMP signalling in the primary cilium. Front Physiol. 2023;14(May):1–13. doi:10.3389/fphys.2023.1187134

34. Kashmiry A, Tate R, Rotondo G, Davidson J, Rotondo D. The prostaglandin EP4 receptor is a master regulator of the expression of PGE2 receptors following inflammatory activation in human monocytic cells. Biochim Biophys Acta - Mol Cell Biol Lipids. 2018;1863(10):1297–1304. doi:10.1016/j.bbalip.2018.07.003

35. Gogendeau D, Lemullois M, Le Borgne P, et al. MKS-NPHP module proteins control ciliary shedding at the transition zone. PLoS Biol. 2020;18(3):1–28. doi:10.1371/journal.pbio.3000640

36. Garcia H, Serafin AS, Silbermann F, et al. Agonists of prostaglandin E2 receptors as potential first in class treatment for nephronophthisis and related ciliopathies. Proc Natl Acad Sci U S A. 2022;119(18):1–11. doi:10.1073/pnas.2115960119

37. Biswas S, Bhattacherjee P, Paterson CA. Prostaglandin E2 receptor subtypes, EP1, EP 2, EP3 and EP4 in human and mouse ocular tissues - A comparative immunohistochemical study. Prostaglandins Leukot Essent Fat Acids. 2004;71(5):277–288. doi:10.1016/j.plefa.2004.03.021

38. Bastien L, Sawyer N, Grygorczyk R, Metters KM, Adam M. Cloning, functional expression, and characterization of the human prostaglandin E2 receptor EP2 subtype. J Biol Chem. 1994;269(16):11873–11877. doi:10.1016/s0021-9258(17)32654-6

39. Osborne NN, Li GY, Ji D, et al. Expression of prostaglandin PGE2 Receptors under conditions of aging and Stress and The protective effect of the EP2 agonist butaprost on Retinal Ischemia. Investig Ophthalmol Vis Sci. 2009;50(7):3238–3248. doi:10.1167/iovs.08-3185

40. Stjernschantz J, Selén G, Astin M, Resul B. Microvascular Effects of Selective Prostaglandin Analogues in the Eye with Special Reference to Latanoprost and Glaucoma Treatment. Vol 19.; 2000. doi:10.1016/S1350-9462(00)00003-3

41. Lorenz B, Gyurus P, Preising M, et al. Early-onset severe rod-cone dystrophy in young children with RPE65 mutations. Investig Ophthalmol Vis Sci. 2000;41(9):2735–2742.

42. Schindelin J, Arganda-Carreras I, Frise E, et al. Fiji: An open-source platform for biological-image analysis. Nat Methods. 2012;9(7):676–682. doi:10.1038/nmeth.2019

43. Bankhead P, Loughrey MB, Fernández JA, et al. QuPath: Open source software for digital pathology image analysis. Sci Rep. 2017;7(1):1–7. doi:10.1038/s41598-017-17204-5

44. Mutterer J, Zinck E. Quick-and-clean article figures with FigureJ. J Microsc. 2013;252(1):89–91. doi:10.1111/jmi.12069

