## Supplementary Table and Figures for "Pharmacological cAMP stimulation via prostaglandin receptors rescues ciliary defects in CEP290-deficient human and mouse models"

**Supplementary Table 1**: Primer pairs specific to PTGER_1_, PTGER_2_, PTGER_3_ and PTGER_4_ mRNAs and sizes of the amplified products

| **Gene** | **Forward primer (5’-3’)** | **Reverse primer (5’-3’)** | **Expected size of the PCR product (bp)** |
| --- | --- | --- | --- |
| *PTGER_1_* | CATGGTGGTGTCGTGCA | TGTACACCCAAGGGTCCAG | 149 |
| *PTGER_2_* | CCTCATTCTCCTGGCTATCATG | CTTTCGGGAAGAGGTTTCATTC | 94 |
| *PTGER_3_* | GCTTATGGGGATCATGTGCG | GCTTATGGGGATCATGTGCG | 100 |
| *PTGER_4_* | ATCTTACTCATTGCCACCTC | CTCGCTCCAAACTTGGCTGA | 166 |

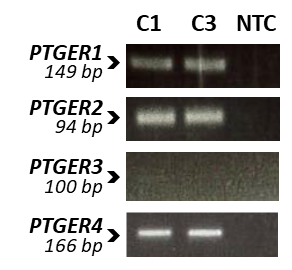

**Figure S1: *PTGER_1_, PTGER_2_, PTGER_3_* and *PTGER_4_* mRNA expression in human primary fibroblasts.** RT-PCR analysis of *PTGER_1_*, *PTGER_2_*, *PTGER_3_*, *PTGER_4_* transcripts in control (C1 & C3) human primary fibroblasts. Images of agarose gel displaying amplicons generated using specific primer pairs for each gene cDNA. A no-template (NTC) reaction was included as a negative control.

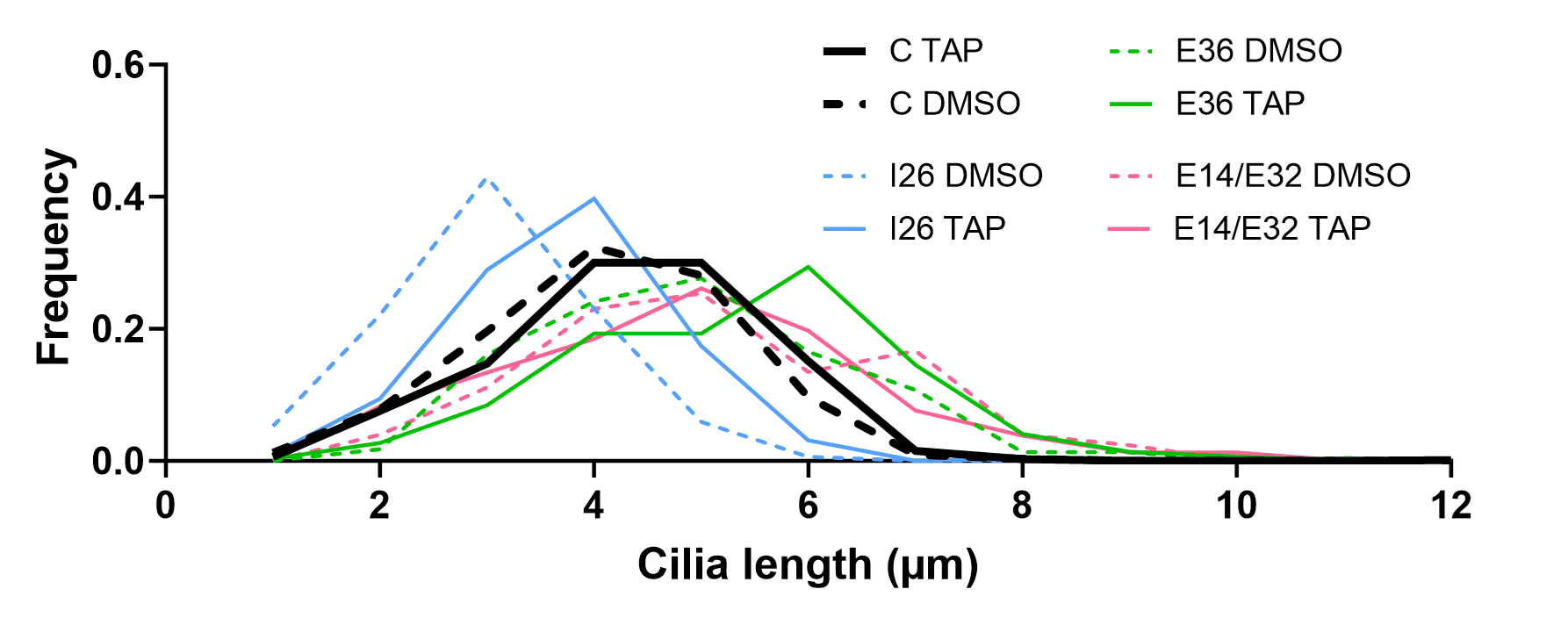

**Figure S2: Distribution of cilia lengths in control and mutant fibroblasts.** This distribution represents cilia lengths in control (C: pool of C1 and C2; bold dark) and mutant (I26, blue; E36, green; E14/E32, pink) fibroblasts under serum starvation after incubation with 0.2µM of Taprenepag in DMSO (solid line) or DMSO alone (dotted line). The cilia lengths were collected from ≥ 3 independent replicates, with a minimum of 300 cells for each condition. ARL13B labeling was used to mark the ciliary membrane, and Gamma-tubulin marked the basal body.

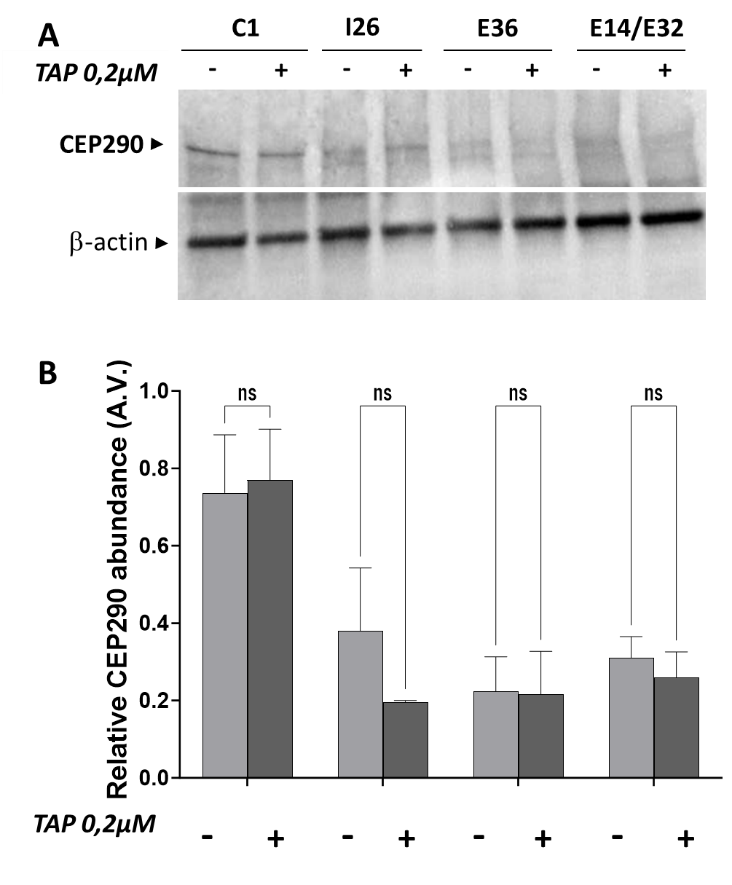

**Figure S3: Effect of Taprenepag on CEP290 Expression in control and mutant fibroblasts.** **A.** Immunodetection of the CEP290 protein in control (C1) and mutant fibroblasts from each genotype (I26, E36, E14/E32) under serum starvation after 24h exposure to DMSO (-) or 0.2µM Taprenepag (+). Beta-actin was used for normalization.

**B.** Quantification of the relative CEP290 abundance in the fibroblasts from controls (C represents the pooled values of C1 and C3) and mutant fibroblasts (I26, E36, E14/E32) under treatment with DMSO (light grey) or 0.2µM Taprenepag in DMSO (dark grey). Beta-actin was used for normalization. Bars represent the mean ± SEM from 3 independent replicates. Statistical test: Two-way ANOVA, with Tukey multiple comparisons. “ns” means no significance.

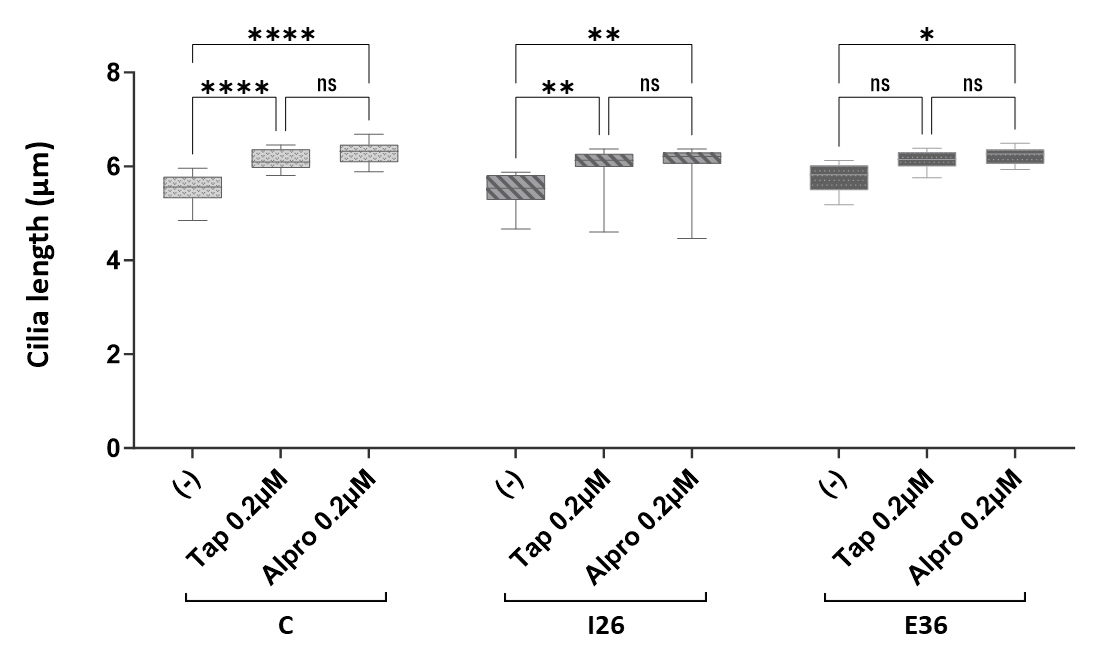

**Figure S4: Comparative effect of Taprenepag and Alprostadil on cilia lengths in control and mutant fibroblasts.** This figure illustrates the average ciliary lengths of control (C: pool of C1 and C3) and mutant (I26, E36) fibroblasts after 24 hours of serum starvation under "high-content" conditions (refer to Material & Methods for details). The cells underwent serum starvation, followed by a 24-hour exposure to either DMSO (-), 0.2µM Taprenepag, or 0.2µM Alprostadil, both diluted in DMSO. Data are presented using a boxplot graphical view, where the line running horizontally within the box represents the median. The upper and lower limits of the box are formed by the 25th (lower quartile Q1) and 75th (upper quartile Q3) percentile values, respectively. The tips of the whiskers extending from the box are the 10th and the 90th percentile, respectively. Data were collected from 3 independent replicates. An ordinary two-way ANOVA was performed with a Tukey's multiple comparison test within each cell line. Significant p-values are denoted as follows: **** for p < 0.0001, ** for p < 0.01, * for p < 0.05, and ns for not significant.

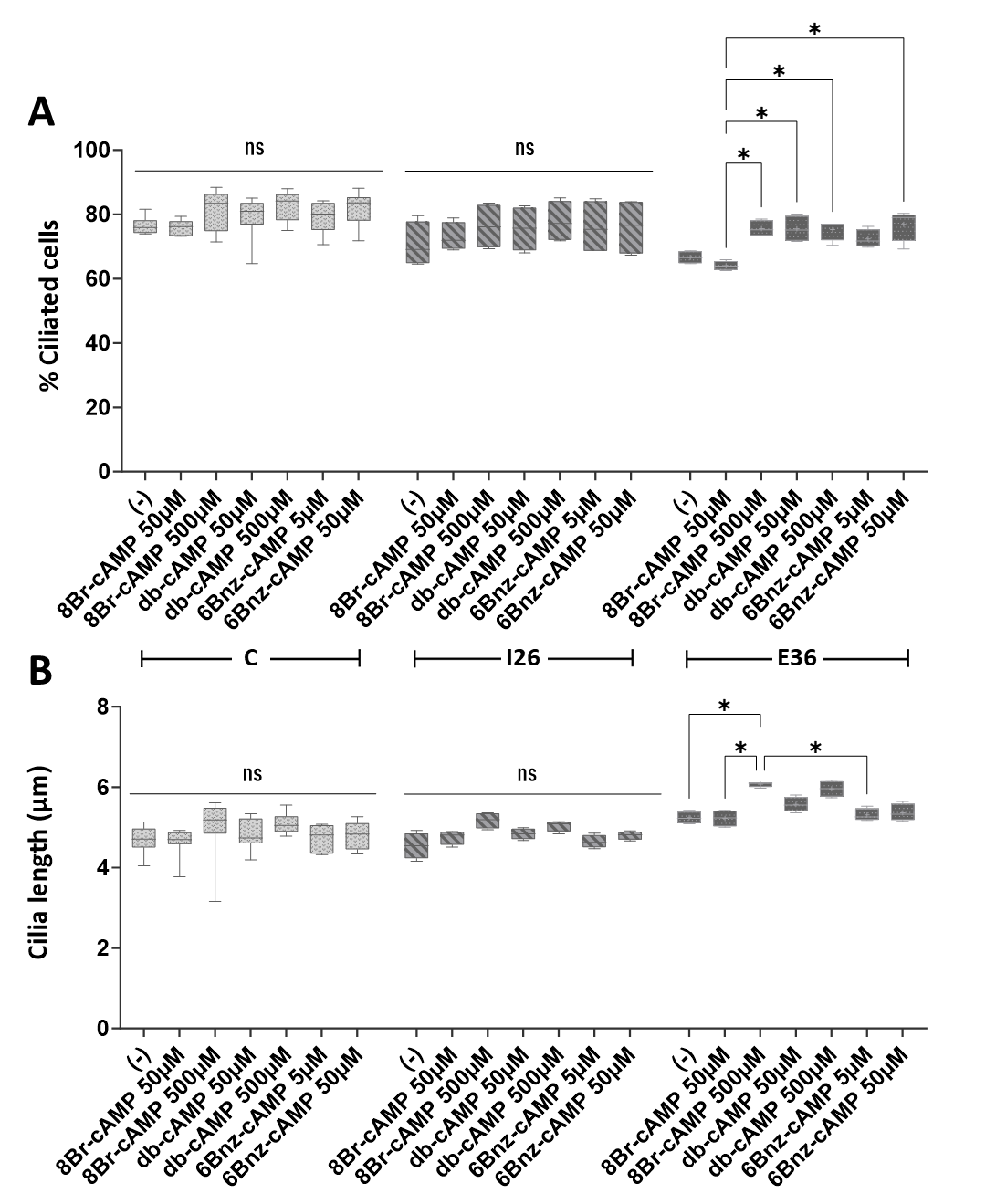

**Figure S5: Effect of cAMP analogs on the cilia formation and length in control and mutant fibroblasts.** Percentage of ciliated cells (**A**) and cilia length average (**B**) as measured by an automatic analysis of serum-starved controls (C) and mutant (I26, E36) fibroblasts either maintained for an additional 48h to serum-free medium only (-) or exposed for 48h exposure to the following cAMP analogs: 8Bromo-cAMP (8Br-cAMP) at 50 and 500µM; dibutyryl-cAMP (db-cAMP) at 50 and 500µM; or 6-benzen-cAMP (6Bnz-cAMP) at 5 and 50µM; all diluted in distilled water. Data are presented using a boxplot graphical view, where the line running horizontally within the box represents the median. The upper and lower limits of the box are formed by the 25th (lower quartile Q1) and 75th (upper quartile Q3) percentile values, respectively. The tips of the whiskers extending from the box are the 10th and the 90th percentile, respectively. Data were collected from 2 independent replicates. An ordinary two-way ANOVA was performed with a Tukey's multiple comparison test within each cell line. P-values <0.05 are indicated by *; ns or no indication = not significant.

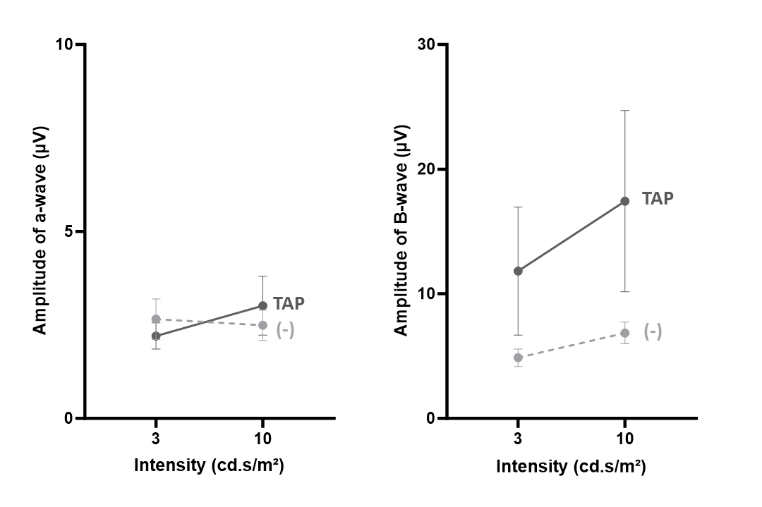

**Figure S6: Taprenepag Effect on *Cep290^del36/del36^* cone response to light stimulation.** Light-adapted (photopic) ERG recordings in P30 *Cep290^del36/del36^* mice after repeated intraperitoneal injections of 18mg/kg Taprenepag (dark grey) or vehicle (dotted light grey). Amplitude of the a-wave (left) and B-wave (right). Bars correspond to the mean ± SEM from 6 mice (left and right eyes). An ordinary two-way ANOVA was performed with Sidak’s multiple comparison test, with no statistically significant observations.
